# The ABCG1 transporter facilitates sesquiterpene accumulation in *Marchantia polymorpha* oil bodies

**DOI:** 10.1101/2025.02.05.636625

**Authors:** Edith C. F. Forestier, Paola Asprilla, Facundo Romani, Ignacy Bonter, Eftychios Frangedakis, Jim Haseloff

## Abstract

*Marchantia polymorpha* oil bodies (OBs) are specialised cell structures housing a diverse array of C15-terpenes, called sesquiterpenes. These compounds are known for their roles as herbivore repellents, yet the enzymes responsible for the biosynthesis of their precursors (C5 isoprenoid units) remain poorly characterized. Discrepancies remain between enzyme localizations suggested by computational predictions and those observed in earlier experimental studies, complicating our understanding of terpene biosynthesis. We investigated the localization of isoprenoid biosynthetic enzymes using translational and transcriptional reporters, coupled with confocal microscopy. Most enzymes localized as predicted (*e.g*., cytosol, chloroplast and the endoplasmic reticulum), and OB cells were identified as the primary sites of terpene biosynthesis.

To explore OBs as potential storage sites for terpenes, we attempted to produce exogenous but easily identifiable compounds in *Marchantia*, such as the diterpene taxadiene and the triterpene β-amyrin. Targeting to OB cells resulted in measurable amounts of these compounds, but their yields remained unaffected by the over-expression of key precursor genes, underscoring challenges in redirecting metabolic flux.

To further investigate terpene accumulation in OBs, we focused on ABCG1, an ABC transporter previously reported to localize at the OB membrane. Overexpression of ABCG1 in OB cells, alongside an exogenous sesquiterpene synthase, only increased the levels of endogenous sesquiterpenes, while CRISPR-mediated disruption of ABCG1 resulted in a dramatic reduction in sesquiterpene accumulation. These findings establish ABCG1 as a critical factor for sesquiterpene retention within OBs and provide new insights into the mechanisms governing terpene metabolism and storage in *Marchantia polymorpha*.

## INTRODUCTION

In recent years, *Marchantia polymorpha* has emerged as a useful model plant system^1^ for studying terpene biosynthesis and accumulation within specialized storage structures known as oil bodies (OBs)^2^. The oil bodies in *Marchantia* are membrane-bound compartments that primarily house terpenes^3^ and bisbibenzyls^4^, a class of phenylpropanoid derivatives; the mixture confers anti-feedant properties to protect against herbivores^5^. Terpenes are a diverse class of natural products derived from the condensation of isoprene (C_5_H_8_) units^6^, found across many organisms but predominantly in the plant kingdom. Beyond deterring herbivores as observed in *Marchantia*, they serve other vital roles in plants, such as preventing fungal infections or attracting pollinators. Additionally, a broad range of bioactive properties makes them valuable in medicine, agriculture, food and other industrial applications.

The sequestration of bisbibenzyls and terpenes into OBs is critical for protecting *Marchantia* from the potential toxicity of these metabolites when released into other tissues. The unique compartmentalization offered by OBs presents a promising avenue for metabolic engineering, as it provides a natural reservoir for accumulating metabolites that might otherwise be harmful to the plant.

Recent developmental studies have provided significant insights into the mechanisms governing OB cell fate determination and formation, highlighting key transcription factors (TFs) involved in these processes. Specific TFs, such as ERF13^7^, C1HDZ^5^, TGA^8^ and MYB2^9,10^ have been identified as crucial regulators of OB cell differentiation and maturation^11^. These discoveries have expanded our understanding of how OB cells develop and contribute to the plant’s metabolic capacity.

Manipulating these TFs has further revealed the delicate balance between OB cell proliferation and plant fitness. For instance, CRISPR-mediated mutation of *TGA*^8^ or gain-of-function mutants of *ERF13* led to a dramatic increase in OB cell numbers per plant^7^. However, these alterations were accompanied by reduced growth rates or morphological defects, suggesting that OB cell number is tightly regulated in *Marchantia* to avoid compromising overall plant health and fitness, or that these TFs have pleiotropic effects that influence additional developmental processes.

Early studies on *Marchantia’s* terpene biosynthetic pathway, which were based on immunolocalization of isoprenoid enzymes, hypothesized that key enzymatic steps might occur at the oil body (OB) membrane^3^. Today, with the availability of whole-genome sequencing and detailed gene annotations informed by sequence homology with other species, our understanding of the early steps of isoprenoid biosynthesis has significantly improved^12–14^ (Table 1 and S1). The mevalonate (MVA) and methylerythritol phosphate (MEP)^15–17^ pathways are the two primary metabolic routes responsible for producing the precursors — isopentenyl pyrophosphate (IPP) and dimethylallyl pyrophosphate (DMAPP) — for various classes of terpenes (Table 1). The MVA pathway, comprising seven enzymatic steps^18^ and typically operating in the cytosol^19^, endoplasmic reticulum^20^ (ER) and peroxysome^21^, generally supplies precursors to sesquiterpenes and triterpenes synthesis^22,23^, while the MEP pathway, with eight enzymatic steps and primarily localized in plastids^24^, commonly provides precursors for monoterpenes and diterpenes^22,23^ (Table 1 and S1). In *Marchantia*, sesquiterpenes are the dominant terpene class found in OBs^2^, produced by several microbial- and fungal-type terpene synthases^25^. Additionally, *Marchantia* produces the monoterpene limonene, likely synthesized through the cis-prenyl precursor neryl pyrophosphate (NPP) rather than the trans form geranyl pyrophosphate (GPP), as demonstrated by Kumar et al^26^. While cell-type-specific metabolic analyses revealed that only sesquiterpenes are stored inside the OB compartment^2^, plants defective in OB cell differentiation show reduced levels of both sesquiterpenes and monoterpenes^5^, making it unclear whether limonene is specifically stored within OBs. Examination of genome data^12–14,27^ along with computational predictive tools for enzyme localization such as DeepLoc 2.1^28^, enabled us to identify putative precursor genes involved in prenyl phosphate synthesis for terpene production (Table 1). However, discrepancies remain between earlier theories suggesting that terpene biosynthetic enzymes are localized to OB membranes^3^ and more recent predictions that place MVA enzymes in the cytosol, ER and peroxisome, and MEP enzymes in plastids (Table 1). These discrepancies prompted us to re-examine the localization of key enzymes using modern techniques such as confocal microscopy combined with translational reporters^7^. This approach allowed us to identify the specific cells expressing these genes and determine the subcellular localization of their corresponding proteins.

**Table 1.**
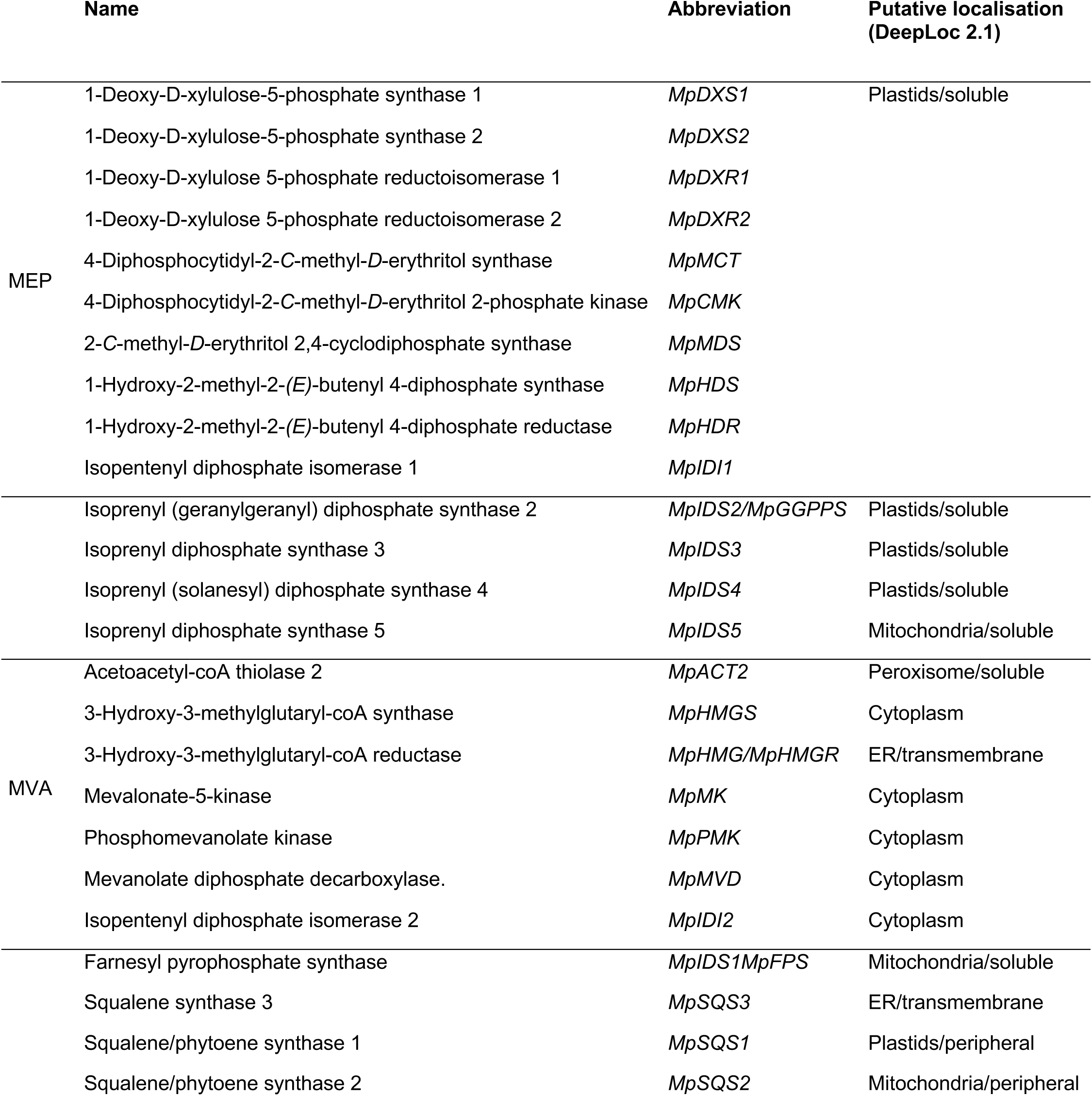
Overview of MVA, MEP, and terpene scaffold biosynthetic genes in *Marchantia polymorpha*. This table lists key genes studied or referenced in this work, including those involved in the methylerythritol phosphate (MEP) and mevalonate (MVA) pathways, as well as enzymes responsible for synthesizing terpene scaffolds for different sub-classes. Abbreviations correspond to gene names, and their putative protein localizations were predicted using DeepLoc 2.1. Detailed annotations and functional descriptions for each gene are provided in Table S1.

Building on our findings regarding terpene precursor gene expression and localization, we then explored the potential for metabolic engineering in *Marchantia polymorpha*’s OBs. Specifically, we aimed to produce two valuable terpenes within OBs: the diterpene taxadiene^29^, a precursor to the valuable anti-cancer compound TaxolⓇ^30,31^, and the triterpene β-amyrin^32^, a precursor to compounds such as glycyrrhizin^33^, a sweet-tasting constituent of liquorice. These compounds were selected due to their distinct chemical signatures, which facilitate easy detection in chromatographic analyses given the minimal background of these terpene sub-classes in *Marchantia*. To further evaluate the metabolic capacity of *Marchantia* and investigate the transport of sesquiterpenes, we also examined the production of the sesquiterpene amorpha-4,11-diene^34^ alongside ABCG1, an ATP-binding cassette (ABC) transporter highly and exclusively expressed in OB cells^35^. Prior studies have demonstrated its clear localization to the OB membrane using a translational reporter^7^, suggesting a potential role in metabolite transport. This study provides new insights into the involvement of ABCG1 in terpene accumulation within OBs and contributes to a broader understanding of the partitioning of terpene metabolism in *Marchantia polymorpha*.

## MAIN

### 1. Translational and transcriptional reporters for genes expression in the terpenoid pathway

To determine the expression pattern and subcellular localization of key enzymes involved in *Marchantia*’s terpene biosynthesis, we generated constructs of translational reporters, in which each gene’s native promoter drove its respective coding sequence fused to the fluorescent tag *mVenus*^36^. In cases where translational reporters were unavailable due to technical challenges, transcriptional reporters were used instead to determine cell-type expression only. These consisted of *Marchantia*’s native promoter driving mVenus with a nuclear localisation signal^37^ (mVenus-N7), enabling us to investigate the spatial and temporal expression of key genes. Promoter lengths were selected based on Assay for Transposase-Accessible Chromatin with Sequencing (ATAC-seq)^38^ data from version 6.1 of the Tak accession in the *Marchantia.info* database^13,25^ (Table S2). ATAC-seq identifies regions of open chromatin, which are more accessible to transcription factors and associated with regulatory domains that influence gene expression. Since relaxed chromatin is frequently found near the start of a gene, it serves as a useful indicator for selecting promoter regions likely to drive gene expression in their native cellular contexts. To delineate cell boundaries, each construct also included a fluorescent membrane marker, *mScarlet-LTI6b* fusion^39–41^, driven by the strong, constitutive *ubiquitin-conjugating enzyme E2* gene promoter and 5’UTR from *Marchantia polymorpha* (*_Prom5_UBE2*)^41^.

For the identification of different subcellular compartments, such as chloroplasts/plastids, ER, cytosol, Golgi, and mitochondria, we interpreted our findings by comparing the localization patterns observed in our images to those of previously characterised standard reporters in *Marchantia*^41,42^, though distinguishing between the ER and cytosol remained challenging under the resolution constraints of our experimental setup.

For this study, we selected key genes from the MVA and MEP pathways on the basis of their predicted roles as rate-limiting enzymes, their redundancy, or their involvement in the final steps of the pathway. Additionally, we included genes responsible for the formation of linear scaffold precursors for sesquiterpenes, diterpenes, and triterpenes to capture a more comprehensive view of terpene biosynthesis. Among these, several genes were annotated as isoprenyl diphosphate synthases (*IDS*)^27^ (Table 1). To ensure accurate functional assignments, we compared their protein sequences with *Arabidopsis* homologs and tentatively re-annotated those likely involved in terpene precursor formation for our study.

We first analysed the two annotated *1-deoxy-D-xylulose-5-phosphate synthases* (*MpDXS1 and MpDXS2*) (Table 1), which encode the first committed and rate-limiting enzymes of the MEP pathway^43^. For *MpDXS1*, tagged with mVenus and driven by its native promoter and 5’UTR (*_Prom5_MpDXS1:MpDXS1-mVenus*), mVenus fluorescence was observed in all cell types throughout the plant, with a stronger signal in OB cells (Figure 1A). The protein localized to plastids, displaying a punctate pattern in non-OB cells rather than a homogeneous distribution (Figure 1A). By contrast, *_Prom5_DXS2:DXS2-mVenus* showed a restricted localization pattern, with fluorescence observed in oil body plastids only from Day 0 gemmae to 14-days-old plants (Figure 1B and S1D).

**Figure 1.**
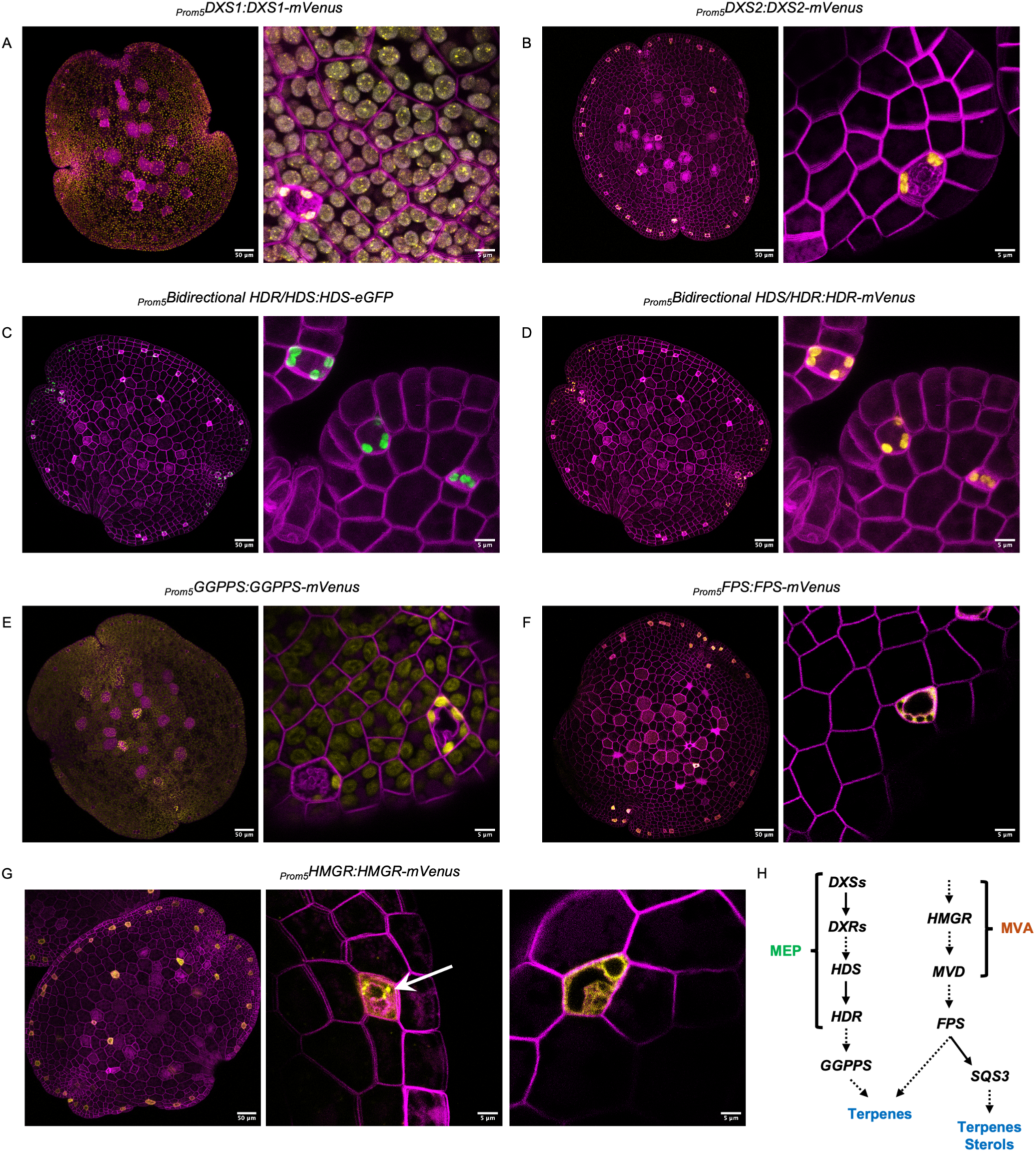
Confocal imaging of translational reporters for key Marchantia polymorpha isoprenoid biosynthetic genes. (A) *_Prom5_DXS1:DXS1-mVenus*, (B) *_Prom5_DXS2:DXS2-mVenus*, (C) *_Prom5_Bidirectional HDR:HDS:HDS-eGFP*, (D) *_Prom5_Bidirectional HDS:HDR:HDR-mVenus*, (E) *_Prom5_GGPPS:GGPPS-mVenus*, (F) *_Prom5_FPS:FPS-mVenus*, and (G) *_Prom5_HMGR:HMGR-mVenus*. Each construct is represented by two images: the full gemmae at day 0 (left panels, scale bar: 50 μm) and a higher magnification view showing subcellular localization (right panels, scale bar: 5 μm). (H) Simplified biosynthetic pathway scheme highlighting enzymatic steps analyzed in this study. The *mVenus* signal (yellow) or eGFP (green) for HDS (C) indicates the subcellular localization of the corresponding enzymes, while *mScarlet* (purple) delineates the cellular boundaries. For the magnified image in (A), chloroplast/plastid autofluorescence (grey) is included to demonstrate sublocalization of DXS1 in the plastids. Panel (G) includes one whole-gemma image and two high magnification images to illustrate the localization of HMGR-mVenus to the ER and Golgi apparatus. White arrow shows the localizations to Golgi.

We examined the final step of the MEP pathway by generating a translational reporter for *(E)-4-hydroxy-3-methylbut-2-en-1-yl diphosphate reductase* (*MpHDR*)^44,45^ (Table 1). We also generated a translational reporter for *(E)-4-hydroxy-3-methylbut-2-enyl diphosphate synthase* (*MpHDS*)^44,46^ (Table 1). *MpHDS* is adjacent to *MpHDR* and oriented in the opposite direction on the same chromosome, suggesting they share a bidirectional promoter (Figure S2A). Previous studies have highlighted the potentially toxic nature of the product of HDS^47,48^, which is detoxified in the subsequent enzymatic step by HDR^47,48^, making it particularly interesting to study both enzymes together. To our knowledge, this is the first report suggesting that *HDS* and *HDR* genes may be regulated by a shared promoter region. Initial experiments revealed that the 2 kb region upstream of *MpHDR* was insufficient to drive expression, whereas the 1.9 kb region upstream of *MpHDS* was adequate (Figures S2B). Expression of a construct driving *MpHDS-eGFP* and *MpHDR-mVenus* bidirectionally (Figure S2C) demonstrated that, like other MEP enzymes, both proteins localized to plastids (Figure 1C and 1D). Their expression was stronger, if not exclusive, to OB cells in Day 0 gemmae (Figure 1C and 1D), before becoming more widespread across tissues at later developmental stages (Figure S1A and S1B).

Finally, we analysed the sub-cellular localisation and cell-type expression of the enzyme forming the linear diterpene precursor geranylgeranyl pyrophosphate (GGPP)^49,50^. Among the IDS candidates^27^, IDS2 and IDS3 share 68% and 62% protein homology, respectively, with GERANYLGERANYL PYROPHOSPHATE SYNTHASE 11 (GGPPS11) from *Arabidopsis thaliana*, the most highly expressed and functionally dominant GGPPS in *Arabidopsis*^51^, which operates as a homodimer^52^. However, transcriptomic data revealed that *IDS3* exhibited consistently low expression across all tissues and developmental stages^12–14^. The other IDS candidates, IDS4 and IDS5, share 71% and 57% protein homology with SOLANESYL PYROPHOSPHATE SYNTHASE and GERANYL PYROPHOSPHATE SYNTHASE, respectively, suggesting potential roles in ubiquinone biosynthesis^53^ (IDS4) and trans-monoterpene precursor formation (IDS5). IDS2 emerged as the strongest candidate for the primary GGPPS in *Marchantia*, given its higher sequence homology and stronger expression. We therefore re-annotated it as MpGGPPS (Table 1 and S1). The translational fusion to *MpGGPPS* showed localization to plastids in all cell types, with particularly strong expression in OB cells (Figure 1E).

For the MVA pathway and sesquiterpene synthesis, we examined the rate-limiting enzyme 3-HYDROXY-3-METHYLGLUTARYL-COA REDUCTASE (MpHMGR)^54^ (Table 1). Expression of *_Prom5_MpHMGR:MpHMGR-mVenus* revealed a restricted localisation to oil body cells in gemmae (Figure 1G) and 14-day-old plants (Figure S1D). The enzyme appeared to display a dual localization pattern: a punctate distribution around the OB, indicative of Golgi, and a web-like pattern surrounding the nucleus, consistent with ER localization (Figure 1G). As a key enzyme downstream the MVA pathway, we analysed FARNESYL PYROPHOSPHATE SYNTHASE (MpFPS) (Table 1), which forms farnesyl pyrophosphate (FPP)^55^, the linear precursor to all sesquiterpenes. Annotated as IDS1^27^ in the Marchantia.info database, MpFPS shares 66% sequence similarity with both FPS homologs from *Arabidopsis thaliana*. Expression analysis revealed that MpFPS is specifically expressed in OB cells, persisting even after 14 days (Figure 1F and S1E). Contrary to the prediction of mitochondrial localization by DeepLoc 2.1^28^ (Table 1), the enzyme appeared predominantly cytosolic, as indicated by its uniform signal surrounding the OB and delineating the plastids (Figure 1F).

We further investigated the expression pattern^56^ of the promoters driving the genes encoding *1-deoxy-D-xylulose 5-phosphate reductoisomerases*^57,58^ (*MpDXR1* and *MpDXR2*) and *mevalonate diphosphate decarboxylase* (MpMVD)^59^ using transcriptional reporters. The corresponding enzymes, MpDXRs and MpMVD, catalyze the second enzymatic step of the MEP pathway and the last enzymatic step of the MVA pathway, respectively (Table 1). *_Prom5_MpDXR1:mVenus-N7* showed ubiquitous expression (Figure 2A), whereas *_Prom5_DXR2:mVenus-N7* showed specificity to OB cells in Day 0 gemma (Figure 2B). Similarly, *_Prom5_MpMVD:mVenus-N7* exhibited specificity to OB cells, even in 14-day-old thalli (Figure 2C).

**Figure 2.**
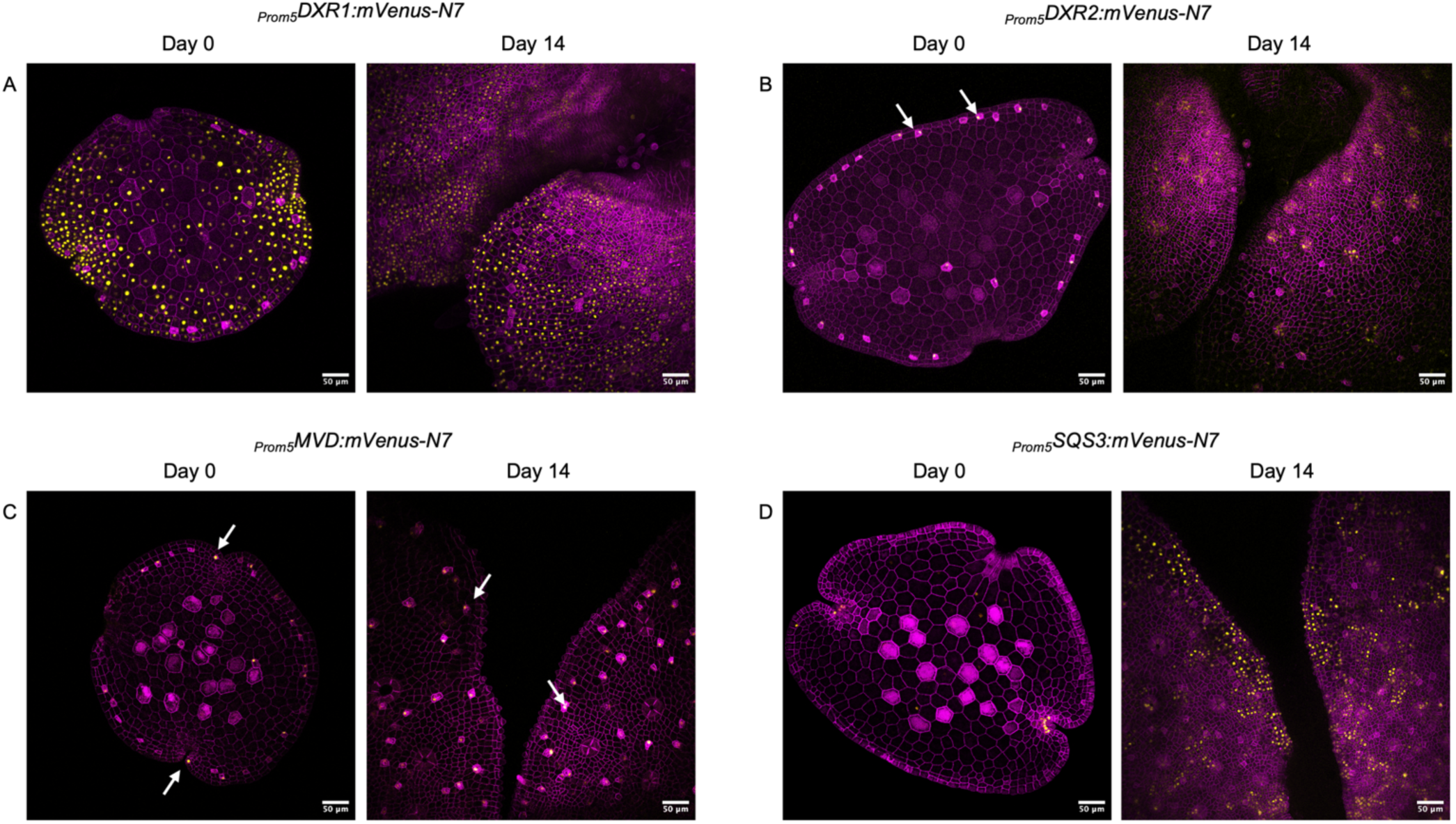
Confocal imaging of transcriptional reporters for additional isoprenoid biosynthetic genes in *Marchantia polymorpha.* (A) *_Prom5_DXR1:mVenus-N7*, (B) *_Prom5_DXR2:mVenus-N7*, (C) *_Prom5_MVD:mVenus-N7*, and (D) *_Prom5_SQS3:mVenus-N7*. The mVenus signal (yellow) localized to the nucleus indicates the activation of these promoters in specific cell types, while mScarlet fluorescence (purple) delineates cellular boundaries. Each construct is represented by two panels: day 0 gemmae (left, scale bar: 50 μm) and the meristem area of 14-day-old plants (right, scale bar: 50 μm). White arrows indicate mVenus signals detected in oil body cells.

Finally, given that the expression of some MVA biosynthetic genes and *MpFPS* appears specific to OB cells, we investigated whether this pattern extends to phytosterol biosynthesis. Phytosterols are essential structural components of plant membranes^60^ and are initially formed through the fusion of two FPP molecules to produce squalene, a reaction catalyzed by SQUALENE SYNTHASE^61^ (SQS). We therefore examined the transcriptional reporter of the *MpSQS* promoter. Among the three squalene synthase-like genes annotated in the *Marchantia* genome (*MpSQS1-3*) (Table 1 and S1), we selected *MpSQS3* based on its higher protein sequence homology (60%) with *Arabidopsis thaliana* SQUALENE SYNTHASE. In contrast, MpSQS1 showed greater similarity (73%) to *Arabidopsis* PHYTOENE SYNTHASE, suggesting a role in carotenoid biosynthesis, while MpSQS2 shared 52% homology with an uncharacterized terpenoid synthase. Plants expressing the *_Prom5_SQS3:mVenus-N7* constructs show a mVenus signal in meristematic regions in Day 0 gemmae, followed by ubiquitous expression in later developmental stages (Figure 2D). These findings indicate that sterol biosynthesis is not confined to OB cells but occurs broadly throughout the plant.

Overall, our findings provide a comprehensive map of the subcellular localization and expression patterns of key terpene biosynthetic enzymes, offering insights into the spatial organization of biosynthetic intermediates and terpene scaffolds in *Marchantia polymorpha*.

### 2. Exogenous production of taxadiene and β-amyrin in *Marchantia polymorpha* whole plants *vs*. oil body cells

Given the observed expression of terpene precursor enzymes predominantly in OB cells, we aimed to harness the metabolic potential of these specialised structures. As proof of concept, we attempted to produce the diterpene taxadiene^29^, and the triterpene β-amyrin^32^.

We generated constructs to express *Taxus baccata taxadiene synthase* (*TXS*)^62^ or *Talinum paniculatum β-amyrin synthase*^63^ (*β-AS*), driven by either the constitutive *_Prom5_UBE2* or the oil body-specific Mp*R2R3-MYB2* promoter (*_Prom5_MYB2*)^37^, enabling expression throughout the entire plant or specifically within OB cells, respectively. TXS catalyzes the cyclization of the linear diterpene precursor GGPP into taxadiene, while β-AS converts the linear triterpene precursor 2,3-oxidosqualene into β-amyrin.

In plants expressing *_Prom5_UBE2:TXS*, we detected a prominent new peak in hexane extracts (Figure 3A), corresponding to taxadiene based on its mass spectrum (MS), as described in prior publications^64,65^. While no commercial taxadiene standards were available, we identified the compound by its characteristic ion at m/z 122 and the molecular ion [M+] at m/z 272 (Figure S3A). Quantification of taxadiene in four independent transformants expressing either *_Prom5_UBE2:TXS* or *_Prom5_MYB2:TXS* revealed significantly higher levels of taxadiene in whole plants compared to those with expression restricted to OB cells (Figure 3D). Specifically, *_Prom5_UBE2:TXS* lines produced approximately 21 µg of taxadiene per g of fresh weight (FW), whereas *_Prom5_MYB2:TXS* lines produced only 0.7 µg/g FW, dodecane equivalent. The correct targeting of TXS protein in OB cells was confirmed using a *_Prom5_MYB2:TXS-mVenus* fusion construct, which showed plastid localization (Figure 3C).

**Figure 3.**
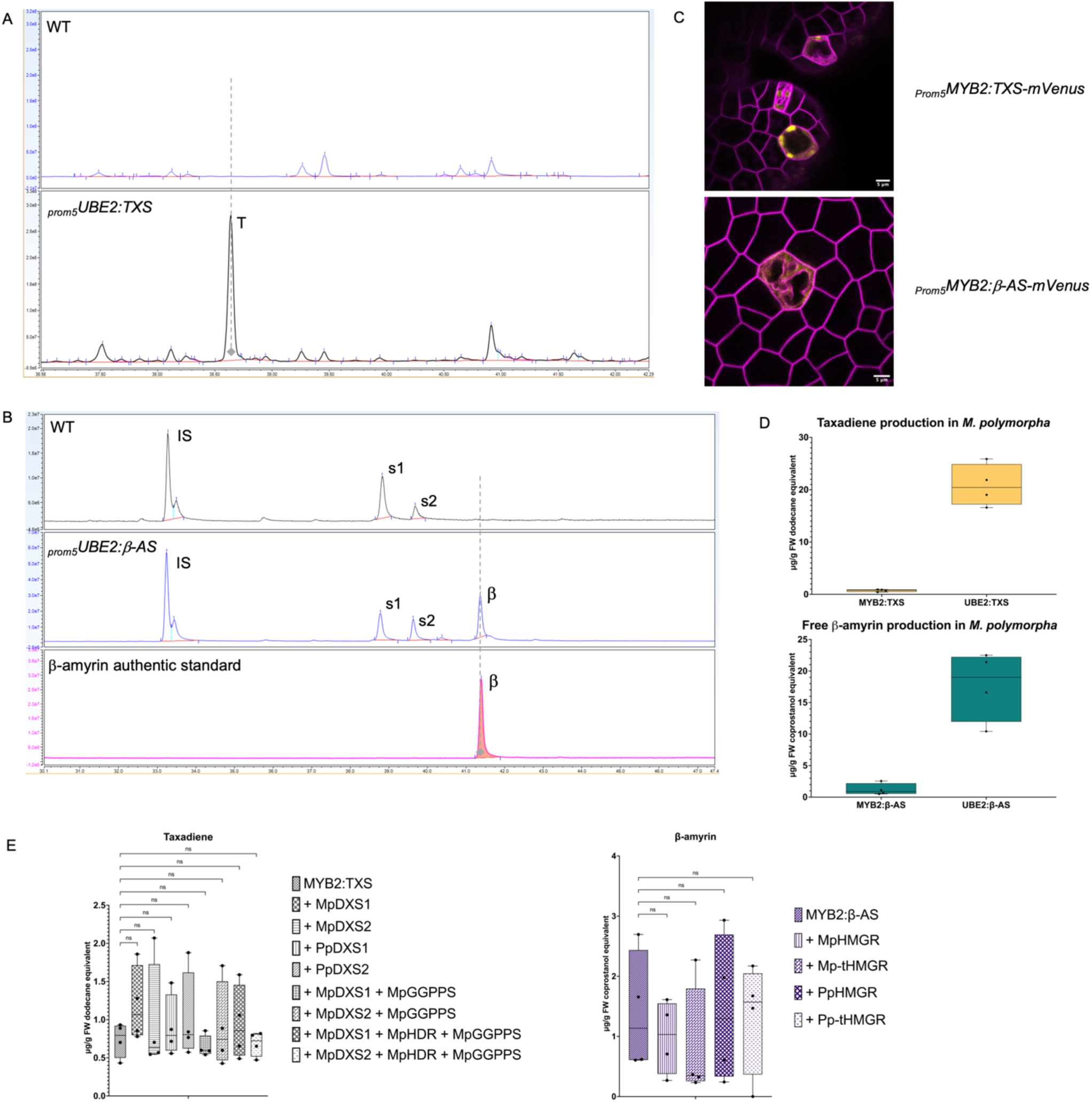
Production of taxadiene and β-amyrin in *Marchantia polymorpha* and subcellular localization of their synthases. (A) Total ion chromatograms (TICs) of terpene extracts from wild-type (WT, upper panel) and transgenic plants expressing *_Prom5_UBE2:TXS* (*taxadiene synthase*, lower panel), highlighting the taxadiene peak (T). (B) TICs of trimethylsilyl-derivatized terpene extracts, showing WT (upper), a transgenic line expressing *_Prom5_UBE2:β-AS* (*β-amyrin synthase,* middle), and an authentic β-amyrin standard (lower). Peaks: IS (internal standard: coprostan-3-ol), s1 (campesterol), s2 (stigmasterol), and β (β-amyrin). (C) Subcellular localization of TXS and β-AS fused to mVenus under the *_Prom5_MYB2* promoter, showing plastid localization for TXS (upper panel) and ER or cytosol localization for β-AS (lower panel). mScarlet fluorescence (purple) delineates cellular boundaries. Scale bar: 5 μm. (D) Quantification of taxadiene (upper graph) and β-amyrin (lower graph) in plants expressing *_Prom5_UBE2:TXS* and *_Prom5_UBE2:β-AS* respectively (whole plant expression) or *_Prom5_MYB2:TXS* and *_Prom5_MYB2:β-AS* respectively (oil body cell-specific expression). Data represent four independent lines per condition. (E) Quantification of taxadiene (left) and β-amyrin (right) in transgenic lines over-expressing precursor genes. For taxadiene, combinations included *MpDXS1*, *MpDXS2*, *PpDXS1*, *PpDXS2*, *MpHDR*, and *MpGGPPS*. For β-amyrin, *MpHMGR*, *Mp-tHMGR*, *PpHMGR*, and *Pp-tHMGR* were tested. Box plots represent data points from independent transformants. One-way ANOVA with Dunnett’s multiple comparison tests (*p* < 0.05) showed no significant differences across gene combinations.

Similarly, plants expressing *_Prom5_UBE2:β-AS* produced a new compound, identified as β-amyrin through its MS profile and comparison with an authentic standard (Figure 3B and S3B). The *_Prom5_UBE2:β-AS* lines produced an average of 56 µg of β-amyrin per g of fresh weight (coprostanol equivalent), whereas lines expressing the enzyme specifically in OB cells yielded an average of 1.4 µg/g FW (Figure 3D). In MYB2-driven *β-AS-mVenus* lines, the protein localized specifically in the cytosol of the OB cells (Figure 3C).

To investigate whether precursor flux limitations were constraining the production of taxadiene and β-amyrin in oil bodies, we co-expressed *TXS* and *β-AS* with key rate-limiting enzymes of the MEP and MVA pathways (*i.e DXS* and *HMGR*, respectively), as our work demonstrated that many of these enzymes appeared to be highly or specifically expressed to OB cells.

For taxadiene production via the MEP pathway, we co-expressed *MpDXS1* or *MpDXS2* with *TXS*. To mitigate potential homology-dependent gene silencing associated with overexpressing native *Marchantia* genes^66^, we also expressed the homologous *PpDXS1* and *PpDXS2* genes from *Physcomitrium patens,* which have recently been re-annotated as *PpDXS1A* and *PpDXS1D*^67^. To further enhance precursor flux, we introduced *MpHDR* and *MpGGPPS*, two key enzymes previously shown to increase GGPP flux for diterpene production in *Nicotiana benthamiana*^68,69^. For β-amyrin production through the MVA pathway^70^, we overexpressed the *HMGR* genes from *Marchantia* (*MpHMGR*) and *Physcomitrium* (*PpHMGR*), alongside truncated versions designed to remove their negative feedback regulatory domain^71,72^, thereby increasing the pool of terpene precursors and allowing unrestrained production of β-amyrin. These truncated variants (*Mp-tHMGR* and *Pp-tHMGR*) lack the first 148 and 146 amino acids, respectively. All genes were expressed under the MYB2 promoter to specifically target OB cells.

Quantitative analysis of four independent transformants showed no significant increase in taxadiene or β-amyrin levels upon over-expression of these precursor supply genes (Figure 3E). We confirmed the correct targeting to OB cells and subcellular localization of *Marchantia DXS*, *HDR*, *GGPPS*, *HMGR*, and *tHMGR* by fusing these genes with mVenus under the control of the MYB2 promoter (Figure S4). DXSs, HDR, and GGPPS enzymes specifically localized to plastids (Figure S4A, S4B, S4C and S4D). However, despite co-localization with TXS, this did not result in increased taxadiene levels. Over-expression of enzymes for precursor biosynthesis in the OB cells didn’t increase either the level of main endogenous sesquiterpenes (Dataset 1 and Dataset 2), which were tentatively identified using Kovats retention indices^73^ (Figure S5 and Table S3). While tHMGR-mVenus appeared to localize in the ER and Golgi apparatus of OB cells (Figure S4E), the native version driven by the MYB2 promoter remained undetectable. To address this, we re-cloned *MpHMGR* under the 2×35S promoter and successfully detected the protein, which accumulated exclusively in the ER of OB cells (Figure S4F). Interestingly, when the truncated version (tHMGR) was expressed under the 2×35S promoter, the fusion protein showed expression across all cells (Figure S4G), suggesting that the negative feedback domain removed in this construct may play a role in restricting localization to OB cells. Overall, these results demonstrate that over-expression of selected rate-limiting enzymes was insufficient to significantly enhance yields of exogenous terpenes in *Marchantia* OB cells, suggesting that additional regulatory mechanisms may play critical roles in terpene biosynthesis and accumulation in these specialized cells.

### 3. Accumulation of terpenes in oil bodies may require a specific transporter

Despite the high specificity and/or strong expression of terpene biosynthetic pathway genes in OB cells, the over-expression of key enzymes did not lead to increased levels of exogenous terpenes. Moreover, these compounds may not naturally accumulate in OBs: taxadiene may primarily localize to plastids, while β-amyrin could accumulate in the cytosol. We hypothesized that precursor flux in *Marchantia* predominantly supplies sesquiterpene biosynthesis and that a specific transporter may be required for their accumulation inside OBs. Previous studies demonstrated high and exclusive expression of the *ABCG1* gene in oil body cells^35^, with its protein localized to the OB membrane when expressed under its native promoter, and to the plasma membrane of other cells when driven by a non-OB-specific promoter^7^.

To investigate whether ABCG1 could facilitate the accumulation of any sesquiterpenes in *Marchantia*’s OBs, we expressed *Artemisia annua amorphadiene synthase* (*AMS*), a sesquiterpene synthase that converts FPP into amorpha-4,11-diene^34^. Cyclisation of FPP into this sesquiterpene likely occurs via the bisabolyl cation^74^, similar to the endogenous β-chamigrene^75^, therefore we expected this structural similarity to enable amorpha-4,11-diene transport into OBs. We targeted *AMS* expression to either the whole plant (*_Prom5_UBE2*) or specifically in OB cells (*_Prom5_MYB2*). Additionally, we co-expressed *AMS* and *ABCG1* in OB cells, alongside *HMGR*, *DXS2*, and *FPS*, with the FPS enzyme recognized as rate-limiting in sesquiterpene biosynthesis^76^. By introducing ABCG1 and FPS, we aimed to overcome potential transport limitations and to test whether this combination could further increase FPP availability for sesquiterpene production. Amorpha-4,11-diene was detected only in transgenic lines expressing the construct *_Prom5_UBE2:AMS* (Figure S6), with an average yield of 0.75 µg/g FW in six out of eight independent transformants (Figure 4A, Dataset 3). No amorpha-4,11-diene was detected when expression was targeted to OB cells, either with or without precursor supply genes and *ABCG1* (Figure S6). Localization studies of *AMS-mVenus* and *MpFPS-mVenus* driven by the MYB2 promoter showed that both proteins localized to the cytosol of OB cells (Figure 4B).

**Figure 4.**
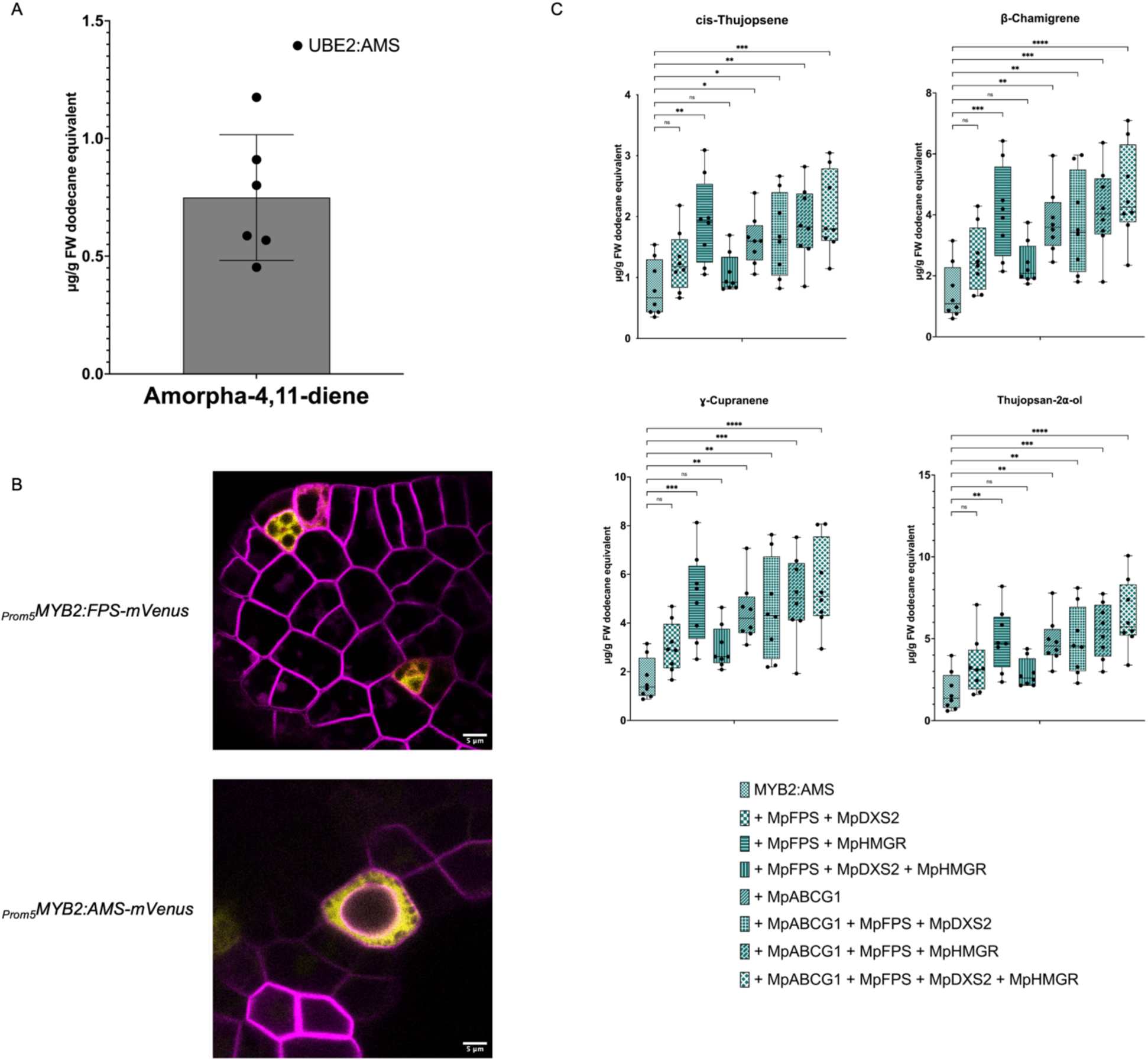
Expression of amorphadiene synthase (*AMS*) in *Marchantia polymorpha* with or without precursor supply genes and the ABCG1 transporter, and its effect on endogenous sesquiterpene levels. (A) Quantification of amorpha-4,11-diene levels in plants expressing *Artemisia annua* AMS under the *UBE2* promoter. The compound was detected in six independent transformants (mean ± SD; n = 6). (B) Subcellular localization of FPS and AMS proteins when expressed under the oil-body-specific *MYB2* promoter. mVenus fluorescence (yellow) indicates protein localization, with signals observed in oil body cells, while mScarlet fluorescence (purple) marks cell boundaries. Scale bars: 5 μm. (C) Quantification of endogenous sesquiterpenes, including cis-thujopsene, β-chamigrene, γ-cuprenene, and thujopsan-2α-ol, in lines expressing *AMS* with or without precursor supply genes (*MpFPS*, *MpDXS2*, and/or *MpHMGR*) and with or without *ABCG1*, under *_Prom5_MYB2*. Box plots represent data points from eight independent transformants (n = 8), with significance determined by one-way ANOVA followed by Dunnett’s multiple comparison test (*p < 0.05*).

We measured endogenous sesquiterpene levels in eight independent transformants for each gene combination. Lines expressing *_Prom5_MYB2:AMS* alone produced relatively low yields, but co-expression of precursor genes significantly increased sesquiterpene levels (Figure 4C, Dataset 3). Among combinations without *ABCG1*, *FPS* + *HMGR* boosted sesquiterpene levels the most, with a 2.8-fold average increase for the major sesquiterpenes compared to a 1.8-fold increase with *FPS* and *DXS2*. The over-expression of *ABCG1* alongside all precursor genes led to the most significant increases in endogenous sesquiterpene levels, with a 3-fold increase in cis-thujopsene, a 3.2-fold increase in β-chamigrene and γ-cuprene, and a 3.6-fold increase in thujopsan-2α-ol (Figure 4C, Dataset 3). The observed increase in endogenous sesquiterpene levels when *HMGR* was expressed under the MYB2 promoter also suggests that HMGR protein is likely localized to OB cells, despite its apparent absence in localization studies using the mVenus tag.

To further validate the transport role of ABCG1, we generated CRISPR lines to disrupt its function. We obtained six independent transformants with early stop codons in the *ABCG1* DNA sequence (Figure 5A). Terpene analysis revealed a dramatic reduction or complete absence of sesquiterpenes in *abcg1* mutants compared to Cas9-only controls (Figures 5B and 5C). Notably, this reduction was specific to sesquiterpenes, as the levels of phytosterols remained unaffected (Figure 5B and 5C). Given the apparent differences in peak abundance observed in the overlaid chromatograms (Figure 5B), we quantified additional compounds alongside neophytadiene and phytol, such as (*Z*)-1,3-phytadiene and (*E*)-1,3-phytadiene, phytol derivatives identified in prior studies^77,78^ (Figure 5B). Quantification revealed no significant changes in these compounds when comparing the five Cas9 controls and six *abcg1* mutant lines (Figure S7). These findings suggest that ABCG1 plays a critical role in the accumulation of endogenous sesquiterpenes while having no effect on other terpene sub-classes or on the accumulation of exogenous sesquiterpenes.

**Figure 5.**
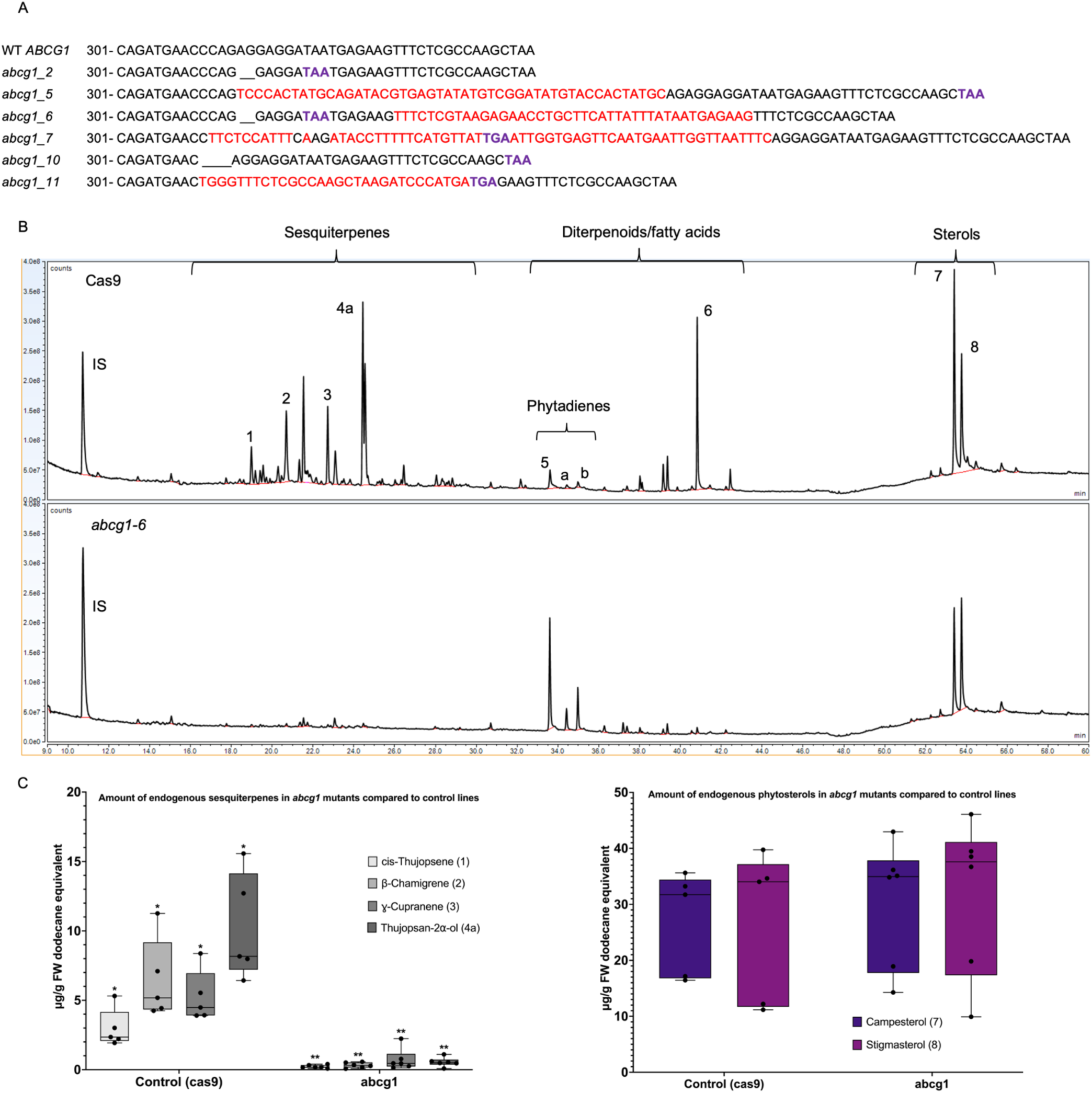
Analysis of endogenous terpene profiles in CRISPR-generated abcg1 mutants compared to Cas9 controls. (A) DNA sequence alignment of the *abcg1* genomic region in wild-type (WT) and CRISPR-generated *abcg1* mutant lines. Dashes indicate deletions, red sequences represent insertions, and purple nucleotides correspond to premature stop codons introduced by the deletions and/or insertions. (B) TICs of terpene extracts from a Cas9 control (upper chromatogram) and abcg1 mutant line (lower chromatogram). Peaks corresponding to quantified compounds are labeled: sesquiterpenes (1: cis-thujopsene, 2: β-chamigrene, 3: γ-cuprenene, 4a: thujopsan-2α-ol), fatty acids/diterpenoids (5: neophytadiene, 6: phytol, a: (*Z*)-1,3-phytadiene, b: (*E*)-1,3-phytadiene), and phytosterols (7: campesterol, 8: stigmasterol). (C) Quantification of endogenous terpene levels in *abcg1* mutants (n=6) compared to Cas9 controls (n=5). The left graph shows the amounts of the four major sesquiterpenes (1 to 4a) labelled in panel B. The right graph displays the levels of phytosterol (7 and 8) labelled in panel B. Box plots display individual data points, with bars representing the median and interquartile range. Statistical significances are denoted by asterisks (one-way ANOVA with Dunnett’s multiple comparison tests (p < 0.05)).

## DISCUSSION

### Mapping of cell-type specificity and sub cellular localisation of isoprenoid biosynthetic enzymes

We used translational and trancriptional reporters to map both the cell-type specificity and subcellular localization of enzymes involved in the early steps of terpene biosynthesis. These findings corroborate earlier transcriptomic analyses in OB-defective plants^5,7^, as well as co-expression network analysis^14^ and single-cell RNA sequencing data^35^, all of which indicate that these enzymes are OB-cell specific.

Enzymatic steps of the MEP pathway and diterpene biosynthesis, including DXS, HDS, HDR, and GGPPS, were localized to chloroplasts/plastids – referred to as chloro-amyloplasts in OB cells by Suire et al^3^ – consistent with their predicted roles in producing isoprenoid precursors for plastid-derived terpenes. In contrast, steps of the MVA pathway and sesquiterpene biosynthesis, represented by HMGR and FPS, were localized to the ER, Golgi, and cytosol, supporting their involvement in cytosolic terpene precursor production. With the exception of FPS, all studied enzymes localized as predicted by DeepLoc2.1^28^. However, our experimental conditions allowed us to localise steady-state accumulation of proteins and did not account for any dynamic behavior due to protein trafficking^79^. Our findings corroborate earlier studies^3^ that reported high or exclusive expression of isoprenoid biosynthetic enzymes in OB cells. Given that OBs are formed through redirection of the secretory pathway^7^, some enzymes like HMGR – which localizes to both the ER and Golgi depending on the OB cell or plant being imaged – may transit to other compartments during OB development, supporting the possibility of their transient presence around the OB membrane^3^.

For each translational or expression pattern reporter, we selected promoter lengths based on a single ATAC-seq^38^ peak, ensuring coverage of the putative core promoter and upstream regulatory regions. In some cases, promoters (including the 5’UTR) were shorter than the typically advised 2.5 kb length^37,41^, balancing the need for regulatory coverage with constraints related to sequence synthesis. For instance, a promoter as short as 970 bp like *_Prom5_GGPPS* was sufficient to drive strong and ubiquitous signal in chloro-amyloplasts. However, in other instances such as *_Prom5_DXR2*, promoter functionality required extending the length from 1.7 kb to 2.7 kb to achieve detectable expression. Altering promoter lengths could include or exclude positive or negative regulatory elements affecting cell-type specificity, therefore further systematic studies would be needed to draw concrete conclusions about the relationship between promoter structure and regulatory outcomes.

Nevertheless, our results confirm the specialized role of OB cells in terpene biosynthesis and highlight the importance of carefully selecting promoters to ensure accurate expression and localization of biosynthetic enzymes.

### Pathway specificity in terpene synthesis: MVA and MEP contributions

Our study suggests that the MVA pathway predominantly provides precursors for sesquiterpene synthesis in *Marchantia polymorpha*, supported by the apparent exclusive expression of HMGR and MVD in OB cells and the significant increases in endogenous sesquiterpene levels observed when *HMGR* and *FPS* were co-expressed. Notably, FPS – also a potential OB-specific enzyme – emerged as a critical limiting step in sesquiterpene biosynthesis, consistent with previous findings in *Nicotiana tabacum*^80^.

The MEP pathway’s contribution to sesquiterpene synthesis cannot be ruled out, given the specific expression of *DXS2* and *DXR2* in OB cells and the fact that a previous study using GC-MS analysis of physically extracted OB content detected sesquiterpenes^2^ as the only sub-class of terpenes. If OBs also store monoterpenes like limonene^26^ or specific diterpenes, their levels remained undetectable or unchanged despite the targeted expression of precursor supply genes. Furthermore, over-expression of key MEP genes and *GGPPS* had no effect on taxadiene yield, despite co-localization of these enzymes with TXS in plastids. This suggests that a significant portion of the precursors is redirected toward sesquiterpene synthesis instead. Supporting this, co-expression of *DXS2* with *FPS*, *HMGR*, and *ABCG1* mildly increased the sesquiterpene levels compared to the same combination of genes without *DXS2*, demonstrating a modest effect of *DXS2* on sesquiterpene yield. Dual contribution from the MVA and MEP pathways in *Marchantia polymorpha* would parallel findings in *Artemisia annua*, where inhibitor assays demonstrated a mixed origin of precursors^81^.

In contrast, *DXS1* and *DXR1* are ubiquitously expressed and likely play broader roles in synthesizing essential metabolites, such as phytyl diphosphate for chlorophyll production. Supporting this, ChIP-seq studies in *Marchantia* have shown that the chloroplast biogenesis regulator GLK binds to the promoter of *DXR1*^82^, further highlighting its role in primary metabolism rather than specialized metabolite production.

The specific localisation of HMGR, MVD, and FPS protein fusions in OB cells raises intriguing questions about how essential metabolites, such as sterols, are produced in non-OB cells. Despite the ubiquitous expression of squalene synthase *SQS3*, which catalyzes the formation of squalene as a precursor to sterols, the source of precursors for sterol synthesis in non-OB tissues remains unclear. Notably, sterol levels in *abcg1* mutants were unaffected, suggesting that OBs are not the primary site of sterol synthesis or storage for the rest of the plant. This is further supported by the observation that sterol levels remain unchanged in plants defective in OB cell differentiation^5^. This raises the possibility that other, as-yet-unannotated MVA enzymes may supply precursors for sterol biosynthesis in non-OB cells. Alternatively, it is plausible that the genes studied here are expressed ubiquitously but accumulate at levels too low to be detected in other cell types with our confocal microscopy studies.

These findings highlight the complexity of precursor flux and compartmentalization in *Marchantia*, accentuating the need to further investigate how primary metabolites synthesis is maintained across tissues.

### Putative role of ABCG1 transporter and implications for metabolic engineering in OB cells

Our study demonstrated that exogenous terpenes, such as amorpha-4,11-diene, taxadiene, and β-amyrin, could be produced throughout the whole plant. However, achieving useful yields within OB cells proved less straightforward, even with the overexpression of key biosynthetic genes. This led us to consider whether transport into OBs might be a limiting factor for the accumulation of exogenous compounds. Supporting this, CRISPR-mediated disruption of *ABCG1* resulted in a sharp reduction in endogenous sesquiterpene levels, demonstrating that ABCG1 plays a crucial role in terpene accumulation. However, the specificity of its transport function remains an open question.

Some ABCG transporters exhibit broad substrate specificity, while others are highly selective. For example, *Nicotiana tabacum* NtNPR1 shows increased ATPase activity when exposed to structurally distinct terpene substrates, including sesquiterpenes like sclareol and capsidiol and the diterpene cembrene^83^, suggesting that it can accommodate chemically diverse terpenes. In contrast, *Arabidopsis thaliana* ABCG29 is a highly specific transporter of *p*-coumaryl alcohol (a lignin precursor)^84^, while several transport assay studies reveal that multiple ABCG transporters exclusively carry the phytohormone abscisic acid (ABA), as reviewed by Do et al^85^. These examples raise the question: does MpABCG1 transport all the sesquiterpenes produced in *Marchantia* OBs, or is it more selective?

Our chromatographic analysis identified over 35 sesquiterpenes in *Marchantia*, though the true number may be even higher, as our GC-MS approach — like previous studies^2^ — primarily detects compounds with limited hydroxylation. If ABCG1 were responsible for transporting all these metabolites into OBs, it would need to accommodate a remarkably high number of substrates. Yet, MpABCG1 did not facilitate the accumulation of exogenous amorpha-4,11-diene, indicating that its function may be more selective than initially expected.

We hypothesize that MpABCG1 could be more exclusive, transporting the sesquiterpene precursor FPP rather than the numerous end-products. To date, no ABCG transporter has been shown to facilitate the movement of a phosphate-containing molecule, with the closest example being *Arabidopsis* ABCC5, an ABCC sub-family transporter that carries inositol hexakisphosphate^86^. However, FPP shares structural features with ABA — not only in its amphiphilic nature but also in its sesquiterpenoid backbone — raising the possibility that an ABCG transporter could accommodate FPP as a substrate.

A serendipitous observation in our study lends further support to the hypothesis that FPP, rather than sesquiterpenes, is transported. A translational reporter of the *fungal terpene synthase-like 2* (*FTPSL2*), an endogenous sesquiterpene synthase characterized by Kumar et al.^26^, was detected within the OB lumen (Figure S8). This finding aligns with earlier theories suggesting that the OB lumen is not merely a storage site but also a catalytic compartment for terpene biosynthesis^3^. However, the conditions under which FTPSL2-mVenus was observed inside OBs remain unclear, as this localization pattern was detected in only one lobe out of four in plants no younger than 14 days old (Figure S8).

If this theory holds true, it implies that precursor flux from both the MVA and MEP pathways is primarily channeled toward FPP production, which is then transported into OBs for conversion by endogenous sesquiterpene synthases. This would explain why co-expression of different *HMGR* versions with *β-AS* did not increase β-amyrin yield: little FPP may remain in the cytosol, limiting its availability for β-amyrin biosynthesis. To determine the precise substrate specificity and transport mechanism of ABCG1, direct biochemical validation of its role in FPP or sesquiterpene transport will be essential. If ABCG1 is indeed specialized in transporting terpene precursors, optimizing taxadiene biosynthesis within OBs would require (1) redirecting GGPPS activity outside plastids, (2) identifying a transporter to mediate GGPP entry into OBs, and (3) engineering TXS for OB localization. Similarly, producing amorpha-4,11-diene or β-amyrin in OBs would necessitate targeting their respective biosynthetic enzymes – and SQS3 in the case of β-amyrin – to this compartment.

We present a preliminary map of terpene biosynthesis in *Marchantia* OB cells (Figure 6). This map includes key steps in primary metabolism, emphasizing the role of certain terpene precursors in vital plant functions, such as sterols, chlorophyll and gibberellin derivative GA ^87^, although the presence of the latter in OB cells has yet to be confirmed (Figure 6).

**Figure 6.**
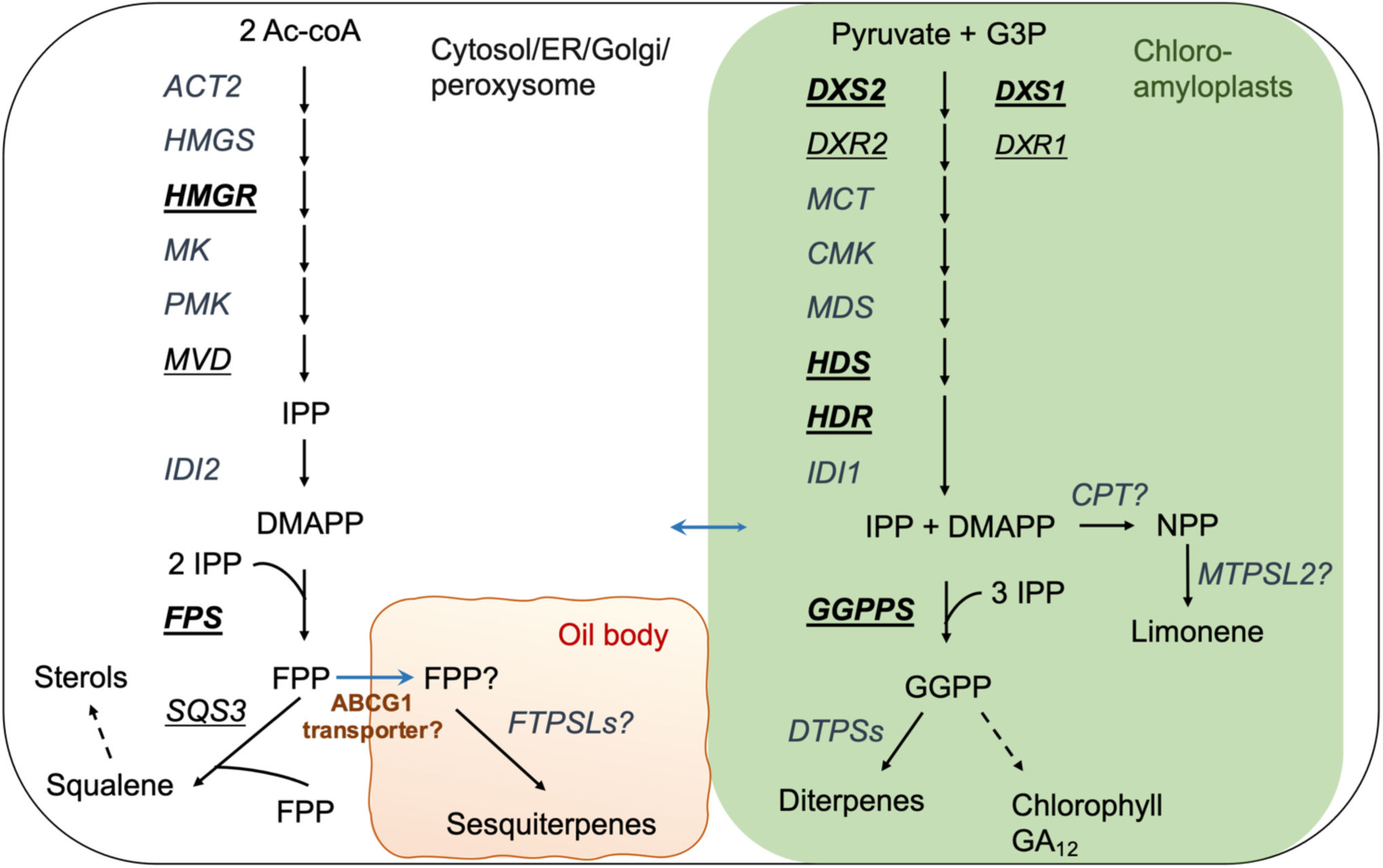
Proposed overview of the isoprenoid precursor pathways in *Marchantia polymorpha* oil body cells based on previous works and findings from this study. Biosynthetic scheme illustrating the mevalonate (MVA) and methylerythritol phosphate (MEP) pathways in *Marchantia polymorpha*. Enzymes shown in bold and underlined were localized using translational reporters confirming their subcellular localization. Enzymes shown in underlined text only were analyzed with promoter expression constructs, indicating activity in oil body cells. Key precursor flows toward sterols, diterpenes, and other metabolites are also indicated. Details of enzymatic steps and annotations can be found in Table 1 and Table S1. Abbreviations: Ac-CoA, acetyl coenzyme A; G3P, glyceraldehyde-3-phosphate; CPT, cis-prenyl transferase; MTPSL2, microbial terpene synthase like 2, functionally characterised in yeast as a limonene synthase (Kumar et al.).

Overall, our work provides a new comprehensive view of terpene synthesis in *Marchantia polymorpha* OB cells by redefining the subcellular localization of key isoprenoid biosynthetic enzymes. Our study also highlights the critical role of ABCG1 in sesquiterpene accumulation and contributes to the broader understanding of metabolic compartmentalization. These findings provide a foundation for engineering terpene biosynthesis with greater precision in specialized plant cells. Beyond *Marchantia*, this work may help uncover similar transport mechanisms in other plant species, offering new strategies for metabolic engineering and plant synthetic biology.

## MATERIAL AND METHODS

### Plant material and growth conditions

*Marchantia polymorpha* subspecies *ruderalis* accessions Cam-1 (male) and Cam-2 (female) were used in this study^88^. Plants were cultivated on solid 0.5× Gamborg B-5 basal medium (#G398, PhytoTech Labs, Lenexa, Kansas, USA) adjusted to pH 5.7–5.8 and solidified with 1.2% (w/v) agar micropropagation grade (#A296, PhytoTech Labs). They were maintained under continuous light at 22°C with a light intensity of 150 μmol/m²/s. For general propagation of the lines, plants were grown in 94 × 16 mm Petri dishes (#633181, Greiner Bio-One, Kremsmünster, Austria). For imaging purposes, plants were cultivated on gridded 65 × 14.5 mm Gosselin™ Petri dishes (#BB64-01, Corning, Corning, NY, USA). Several gemmae, originating from four independent transformants, were grown on these plates and imaged at developmental stages corresponding to Day 3, Day 5, Day 7, and Day 14 after germination.

For gemma cup production and the collection of material for terpene analysis, the media were supplemented with 0.5% (w/v) sucrose. Plants destined for terpene extraction and quantification were grown in 100 × 25 mm Petri dishes (#D943, Phytotech Labs) containing 50 mL of medium.

Spore production was carried out in Microbox micropropagation containers (SacO2) under long-day conditions (16 h light/8 h dark) with light supplemented by far-red illumination, following previously established protocol^41^.

### Protein sequence analysis

Protein sequences of the selected IDS and SQS candidates from *Marchantia polymorpha* were analyzed using NCBI BLASTP (Basic Local Alignment Search Tool for Proteins, https://blast.ncbi.nlm.nih.gov/Blast.cgi) against the *Arabidopsis thaliana* protein database. Sequence similarity was assessed based on percentage identity and coverage to infer functional homology.

### Plasmid construction for over-expression studies and CRISPR-based mutant generation

DNA fragments used to generate mVenus and metabolic constructs were synthesized by Genewiz (Azenta, Burlington, Massachusetts, USA) either as linear fragments or in pUAP1 vector^89^. Linear fragments included overhangs compatible with LguI (#ER1931, ThermoFisher Scientific, Waltham, Massachusetts, USA) cloning into L0 acceptor vectors following the Loop protocol^41^. Linear fragments and sequences synthesized in pUAP1 vectors, contained overhangs enabling direct cloning into L1 or pBy01^37^ vectors using BsaI-HF ^®^v2 enzyme (#R3733S, New England Biolabs, Ipswitch, Massachusetts, USA).

*MpFPS* and *MpHDS* sequences were cloned from the *Marchantia* transcriptome as they didn’t require to remove the internal restriction sites of BsaI and LguI, incompatible with the Loop cloning system. Total RNA was first extracted from *Marchantia* tissue using the RNeasy Plant Mini Kit (#74904, Qiagen, Hilden, Germany) and reverse-transcribed into cDNA with Superscript IV (#18090010, Invitrogen, Waltham, Massachusetts, USA) using random hexamer oligos (#N8080127, Invitrogen) following the protocol from the manufacturer. Target sequences were subsequently amplified from the cDNA using VeriFi® Polymerase Mix (#PB10.43-01, PCR Biosystem, London, UK). PCR products were purified using the QIAquick PCR Purification Kit (#28104, Qiagen) before being used in the cloning steps.

DNA sequences exogenous to *Marchantia* were domesticated when necessary and codon-optimised using the Genewiz codon-optimisation tool prior to ordering. NCBI accession numbers for each exogenous genes are as follows: *TXS* (AY424738), β*-AS* (MG492000), *PpDXS1* (XM_024524883), *PpDXS2* (XM_024533934), *PpHMGR* (XM_024507706) and *AMS* (AY006482). Certain sequences were re-amplified to remove stop codons and introduce the appropriate overhangs for fusion with mVenus or eGFP tags. Primers used in this study were ordered from IDT (Integrated DNA Technology, Coralville, Iowa, USA) (Table S4).

Vectors, promoters, terminators, tags, and pre-assembled cassettes used to generate the constructs were sourced from the OpenPlant toolkit^41^ or adapted from Romani et al^37^. Final plasmids (L2, L3, or pBy01) were sequenced at Plasmidsaurus (Eugene, Oregon, USA) to confirm proper assembly and integration of all transcription units. Details of the sequences and vectors syntax are available in Dataset 5.

The guide RNA sequence used to generate *abcg1* mutants was designed using the CasFinder tool^90^. This guide was ordered as primers with appropriate overhangs, and cloned into a single-guide acceptor vector following the protocol of Sauret-Guëto et al^23^.

### *Agrobacterium*-mediated transformation of *Marchantia* spores and selection of independent transformants

*Marchantia* spores were sterilized and transformed following previously described protocols, with slight modifications. Briefly, a single archegoniophore, dried and stored in silica beads, was mixed in 1.5 mL of a chlorinated solution consisting of one Milton sterilizing tablet (troclosene sodium; Boots, UK) dissolved in 10 mL of sterile water. Spores were released into the solution, filtered through a 40 µm cell strainer (#542040, Greiner Bio-One) and incubated in the sterilizing solution for 30 minutes. After sterilization, spores were harvested by centrifugation, resuspended in sterile water, and spread on Gamborg medium agar plates (without sucrose).

After 5 days of germination, the sporelings were co-cultured with *Agrobacterium tumefaciens* GV3101 carrying the constructs of interest. Transformation of *Agrobacterium* with the constructs was performed using the freeze thaw method^91^. Sporelings and *Agrobacterium* were co-cultured for 2 days in a 4 mL solution of 0.5× Gamborg B-5 plus supplements like previously described^41^, and incubated at 22°C under continuous shaking and light. Following co-culture, sporellings were collected on 70 µm cell strainers (#542070, Greiner Bio-One), rinsed thoroughly with sterile water, and plated on 90 mm Petri dishes containing 0.5× Gamborg B5 medium supplemented with 100 µg/mL cefotaxime (#BIC0111, Apollo Scientific, Bredbury, UK) to eliminate *Agrobacterium* and 20 µg/mL hygromycin (#10687010, Invitrogen) to select for transformants.

After two weeks on the first selective plates, eight transformants were typically transferred to a second plate containing the same selection agents to confirm successful transformation and eliminate residual *Agrobacterium*. For lines expressing fluorescent proteins (mScarlet, mVenus or eGFP), transformants were pre-selected under a Leica stereo microscope (#MDG41, Leica, Wetzlar, Germany) using suitable filter sets to confirm the presence of fluorescence signals. For lines expressing exogenous terpenes, eight plants were randomly chosen, grown for one month on the second selective plates, and subsequently extracted to confirm the presence of the terpene of interest. The four best lines producing the desired terpene were then transferred to sucrose-supplemented plates to quantify endogenous and exogenous terpene levels.

For lines co-expressing *AMS* and precursor supply genes, a single vector carrying up to six transcription units could not be constructed due to cloning or vector size limitations. Attempts to co-transform using two *Agrobacterium* strains carrying distinct plasmids — one with hygromycin B as the selection agent and the other with chlorsulfuron — resulted in an insufficient number of transformants. Therefore, transformants were selected based on hygromycin B resistance for *AMS* (with or without *ABCG1*) on one vector, and the presence of an mScarlet-Lti6b fluorescent signal from the second vector carrying *MpFPS*, *MpHMGR*, and/or *MpDXS2*. All eight independent lines were retained to measure amorphadiene and endogenous terpene level.

To genotype CRISPR-generated *abcg1* mutants and Cas9-only control lines, genomic DNA was extracted using the InnuPrep Plant DNA kit (#845-KS-1060050, IST Innuscreen, Berlin, Germany). For *abcg1* lines, a 500 bp DNA fragment spanning the gRNA target site was amplified using Q5® High-Fidelity DNA Polymerase (#M0491S, New England Biolabs). The resulting PCR products were purified and sequenced by Genewiz to confirm the disruption or not of the *ABCG1* coding sequence. The presence of the Cas9 cassette in control lines was confirmed by amplification with specific primers (listed in Table S4) and subsequent sequencing. A total of 16 lines were genotyped for the abcg1 ones, and eight for the Cas9 controls.

### Laser scanning confocal microscopy

For transgenic lines expressing fluorescent proteins, images were acquired on an upright Leica SP8X confocal microscope equipped with a 460–670 nm supercontinuum white light laser, two continuous wavelength laser lines of 405 nm and 442 nm and a five-channel spectral scanhead (four hybrid detectors and one photomultiplier). Imaging was conducted using either a 25× water immersion objective (Fluotar VISIR 25×/0.95 WATER) for whole-gemmae or meristematic area imaging, or a 40× water immersion objective (HC PL APO CS2 40×/1.10 WATER) for higher magnification images, with an additional digital zoom applied up to a factor of 5× to enhance visualization of subcellular localizations. Excitation laser wavelength and fluorescence emission bandwidth windows were as follows: 515 nm and 525–550 nm (for mVenus); 570 nm and 591-621 nm (for mScarlet); 442 nm and 645–664 nm (for chlorophyll autofluorescence). Each channel was acquired separately using a hybrid detector, with a sequential scan for each channel performed on the Leica LAS X software. To image sequentially the mVenus, eGFP and mScarlet fluorescences on the translational reporter lines carrying the bidirectional construct of *HDS-eGFP* and *HDR-mVenus*, excitation and emission bandwidth windows were adjusted as follow to prevent signal crosstalk: 487 nm and 497-513 nm (for eGFP); 521 nm and 530-550 nm (for mVenus); 570 nm and 588-598 nm (for mScarlet).

Day 0 gemmae were imaged by mounting them on a glass slide with perfluorodecalin^92^ (#130040250, ThermoFisher) and covering them with a glass coverslip. For plants aged 3 to 14 days, a 15 x 16 mm gene frame (#AB0577, ThermoFisher) was placed on the glass slide to prevent compression of the plant material under the coverslip. Whole gemmae and overviews of the meristematic region in older plants were imaged using Z-stack scans, which were processed in Fiji^93^ to generate maximum intensity projections of the Z-stacks. High magnification images were acquired as single scans to better display subcellular features.

### Extraction of terpenes and analysis by gas chromatography-mass spectrometry (GC-MS)

Approximately 200 mg of frozen tissue from 1-month-old *Marchantia polymorpha* plants was extracted with 1 mL of cold methanol containing 5 mM NaCl to quench enzymatic activity, following the method of Kumar et al^26^. To enable quantification, 5 µg of dodecane (#297879, Sigma-Aldrich) was included as an internal standard. The tissue was disrupted in the solvent using a 3 mm stainless steel ball (#2205, Durston, High Wycombe, UK) and homogenized with a TissueLyser II (Qiagen). Samples were subsequently agitated on a Vibrax® shaker (#0002819002, IKA, Staufen, Germany) at 2000 rpm for two hours, and the resulting extracts were centrifuged to remove plant debris. The methanolic phase was then extracted once with hexane to isolate non-polar and medium-polar terpenes from the upper organic layer.

Hexane extracts (200 µL) were analyzed on a Trace 1300 gas chromatograph (ThermoFisher) coupled to an ISQ 700 mass spectrometer (ThermoFisher) with a Zebron CD-5MS column (30 m × 0.25 mm × 0.25 μm; ThermoFisher). A splitless injection (1 µL) was conducted at 230°C, and the GC-MS oven parameters were adapted from Kumar’s work^26^ as follows: the oven temperature was initially set to 70°C and held for 3 min, followed by a ramp of 20°C/min to 90°C, a second ramp of 3°C/min to 180°C, a third ramp of 5°C/min to 240°C, and a final ramp of 20°C/min to 300°C, with a 6-min hold at this temperature. The MS began data acquisition after a 5.5-minute solvent delay, with the transfer line and ion source temperatures set at 250°C and 270°C, respectively. Scanning was conducted in full-scan mode (scan time: 0.17 s) over a mass range of 40–600 atomic mass units. Helium was used as the carrier gas, with a flow rate of 1.2 mL/min for sesquiterpenes in the taxadiene and β-amyrin datasets. For amorphadiene detection, the carrier flow rate was reduced to 0.9 mL/min. Chromatograms were processed using Chromeleon™ software (ThermoFisher), and terpene quantification was performed relative to the internal standard dodecane. Tentative identification of the major sesquiterpenes was achieved by comparing mass spectra to published data and by calculating retention indices using a single sample run with a C8–C40 alkane standard mixture (#40147-U, Superlco, Bellefonte, Pennsylvania, USA) as described by Adams^73^. Note that the endogenous terpene composition in CAM accessions may slightly differ from that reported for other accessions, such as Tak, in previous studies. The authentic standard of amorpha-4,11-diene was provided by Tomasz Czechowski and Ian Graham (University of York).

To achieve more accurate quantification of β-amyrin and sterols, the extraction protocol and GC-MS method were modified. The same amount of frozen material was extracted with cold methanol supplemented with 10 µg of coprostan-3-ol (#C7578, Sigma-Aldrich) as an internal standard. The samples were extracted twice with hexane to obtain a total hexane volume of 1.5 mL, which was then dried under nitrogen flow in a Genevac™ concentrator EZ-2 (#EZ3P-23050-NN0, Genevac Ltd, Ipswich, UK). Extracts were derivatized with 50 µL 1-trimethylsilyl-imidazole (TMS; #92718, Superlco) and resuspended in 300 µL hexane. For GC-MS analysis, 200 µL of the derivatized samples was transferred into vials for direct injection. The method used for sterol/triterpene analysis was as follows: 130°C hold for 2 min; ramp of 30°C/min until 220°C, then 2°C/min until 300°C, hold for 10 min. Solvent delay of 10 minutes before acquiring the MS data. The authentic standard of β-amyrin was provided by James Reed and Anne Osbourne (John Innes Center, Norwich, UK).

For terpene quantification in *abcg1* CRISPR and Cas9 control lines, extractions were performed on genotyped plants from the second selection stage, as loss of the CRISPR genotype was observed during propagation from gemmae. Given this observation, we considered the possibility of chimerism^94^ and extracted as much biomass as was reasonably available from individual plants. This approach aimed to minimize variability, homogenize the material, and ensure accurate results.

Box plots showing individual data points for each quantified terpene were created using PRISM software version 10 (GraphPad, La Jolla, California, USA), and the results of statistical comparisons were integrated into the visualizations. For lines expressing exogenous terpene synthases, statistical analyses were performed using one-way analysis of variance (ANOVA) to assess differences across gene combinations. Homogeneity of variances was evaluated using the Brown-Forsythe and Bartlett’s tests, with Dunnett’s multiple comparison test applied post-hoc to compare each gene combination to the respective control (*_Prom5_MYB2:β-AS*, *TXS*, or *AMS*). The significance threshold (alpha) was set at 0.05. Four biological replicates expressing the constructs were analyzed to quantify both exogenous (β-amyrin and taxadiene) and endogenous compounds (n=4), while eight biological replicates expressing the constructs were analyzed for amorphadiene lines, focusing on the quantification of endogenous terpenes since amorphadiene was not detected (n=8). For comparisons of terpene levels between *abcg1* mutants (n = 6) and Cas9 controls (n = 5), Welch’s t-test was used when variances were unequal, and unpaired t-test was used when variances were equal, as determined by PRISM. Both tests accounted for differences in sample size between the two groups. Asterisks were added to the box plots to indicate statistically significant differences (P<0.05) between *abcg1* mutants and Cas9 control lines, as determined by Welch’s or unpaired t-tests.

## Acknowledgements

This work was funded by the BBSRC NEBP Transition Award BB/W014173/1, with part support from the BBSRC/EPSRC OpenPlant Synthetic Biology Research Centre Grant BB/ L014130/1 to J.H. I.B was funded by a Herchel Smith studentship.

The authors thank Davide Annesse and Connor Tansley (University of Cambridge) for their insightful discussions and support throughout this project. We also thank Anne Osbourn and James Reed (John Innes Centre, Norwich, UK) for providing a β-amyrin standard, as well as Ian Graham and Tomasz Czechowski (University of York) for providing an amorpha-4,11-diene standard.

## Author information

### CONTRIBUTIONS

E.C.F.F. designed the work. F.R. provided DNA parts and early insights. E.C.F.F., P.A., I.B. and E.F. carried out the work. E.C.F.F. wrote the manuscript with input from all authors. All authors approved the final version of the manuscript.

