## Supplementary figures for "The ABCG1 transporter facilitates sesquiterpene accumulation in *Marchantia polymorpha* oil bodies"

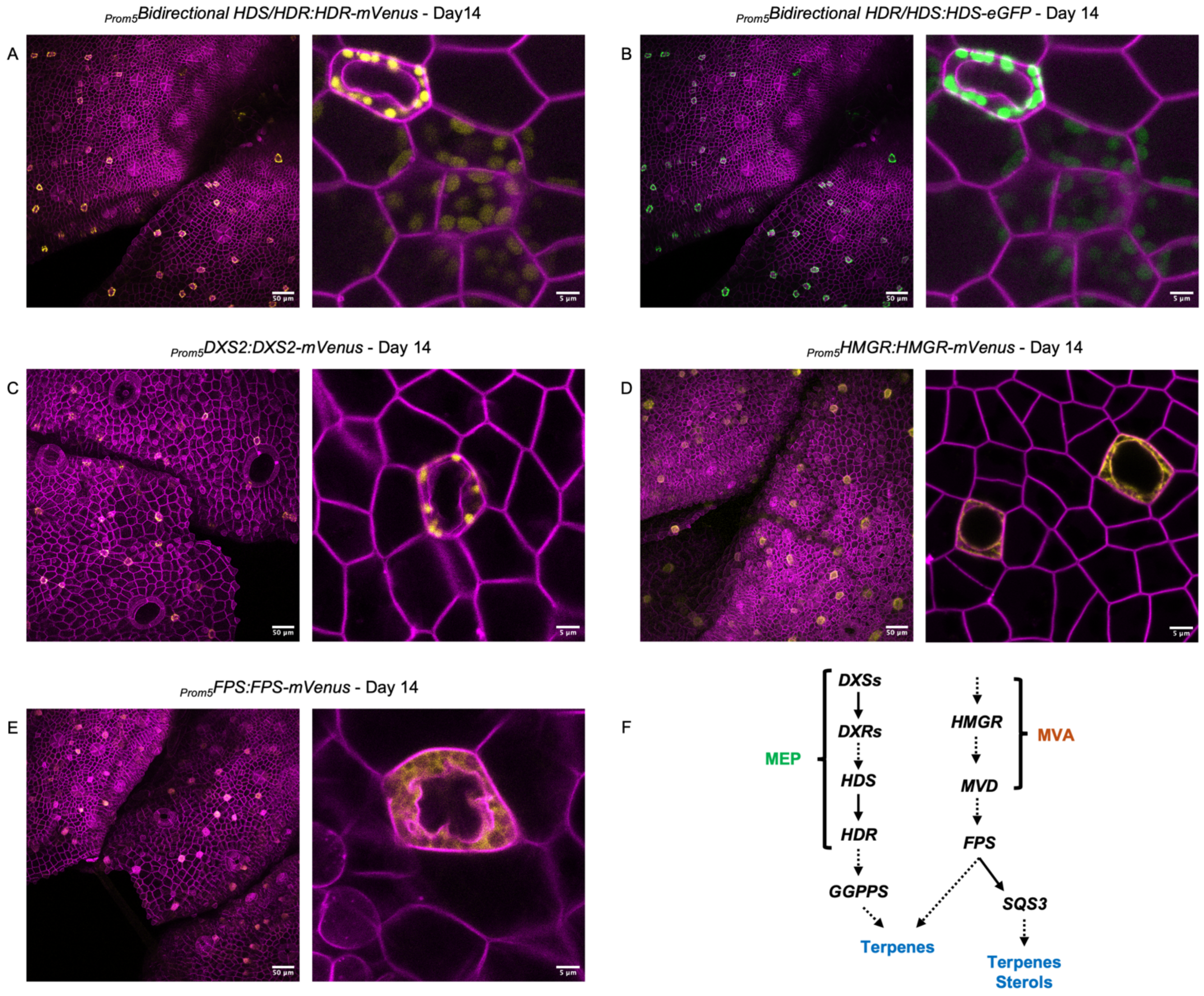


**Figure S1. Confocal imaging of translational reporters for selected *Marchantia polymorpha* isoprenoid biosynthetic genes in the meristem area of 14-day-old plants.** (A) $_{Prom5}$*Bidirectional HDS/HDR:HDR-mVenus*, (B) $_{Prom5}$*Bidirectional HDR/HDS:HDS-eGFP*, (C) $_{Prom5}$*DXS2:DXS2-mVenus*, (D) $_{Prom5}$*HMGR:HMGR-mVenus* and (E) $_{Prom5}$*FPS:FPS-mVenus*. The mVenus (yellow) or eGFP (green) signals indicate the subcellular localization of the respective enzymes, while mScarlet fluorescence (purple) delineates cellular boundaries. Each construct is represented by two panels: the left panels (scale bar: 50 µm) show a broader view of the meristem area, while the right panels (scale bar: 5 µm) provide a zoomed-in view of subcellular localization. (F) Simplified biosynthetic pathway highlighting the enzymatic steps studied using translational and transcriptional reporters studied in this work.

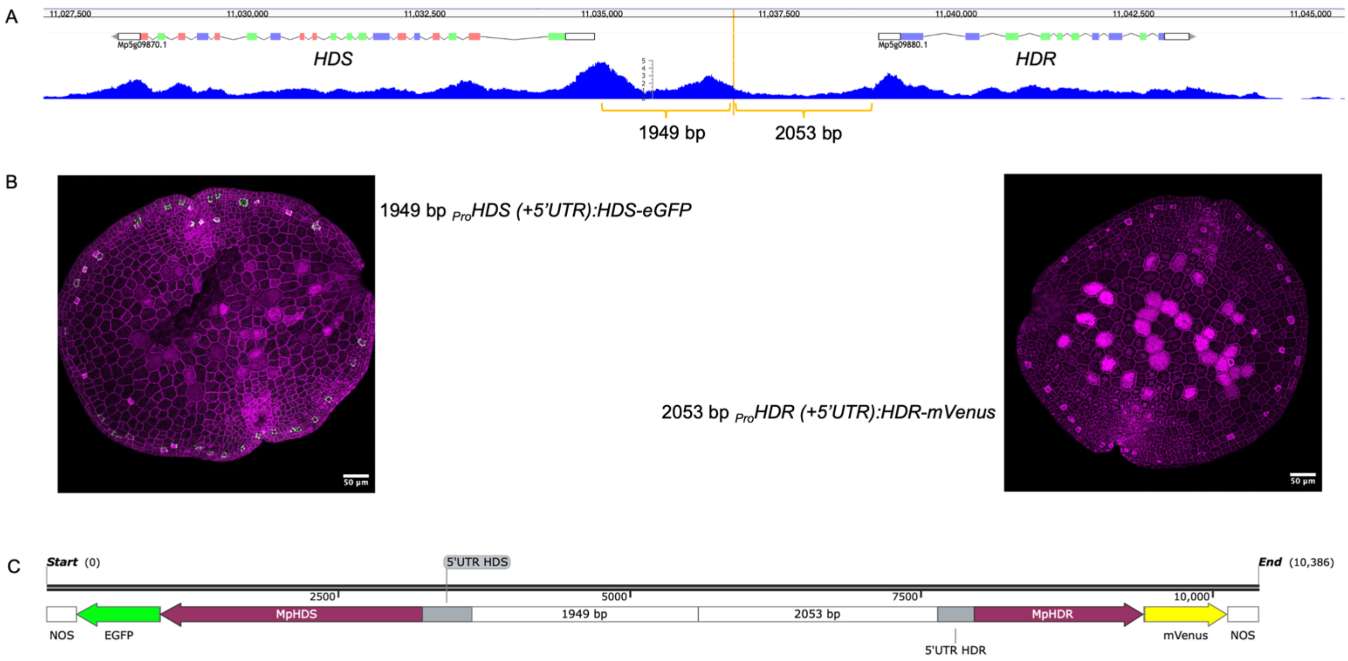


**Figure S2. Bidirectionality of the *HDS-HDR* promoter region and design of the bidirectional translational reporter.**
(A) Gene structure of *HDS* and *HDR*, showing the region between the two genes with ATAC-seq peaks (blue) extracted from the Marchantia.info database (Tak accession, version 6.1). The orange line indicates a boundary splitting the intergenic region to create separate promoters for *HDS* (left) and *HDR* (right). (B) Confocal imaging of translational reporters $_{Prom5}$*HDS:HDS-eGFP* (left) and $_{Prom5}$*HDR:HDR-mVenus* (right) in day 0 gemmae. The mScarlet fluorescence (purple) delineates cell boundaries, while eGFP (green) indicates subcellular localization of HDS. HDR-mVenus fluorescence is not detected. Scale bar: 50 µm. (C) Schematic representation of the bidirectional translational reporter construct designed to drive expression of *HDS* and *HDR,* with mVenus (yellow) and eGFP (green) reporters fused to *HDR* and *HDS*, respectively. The construct design was performed using SnapGene software.

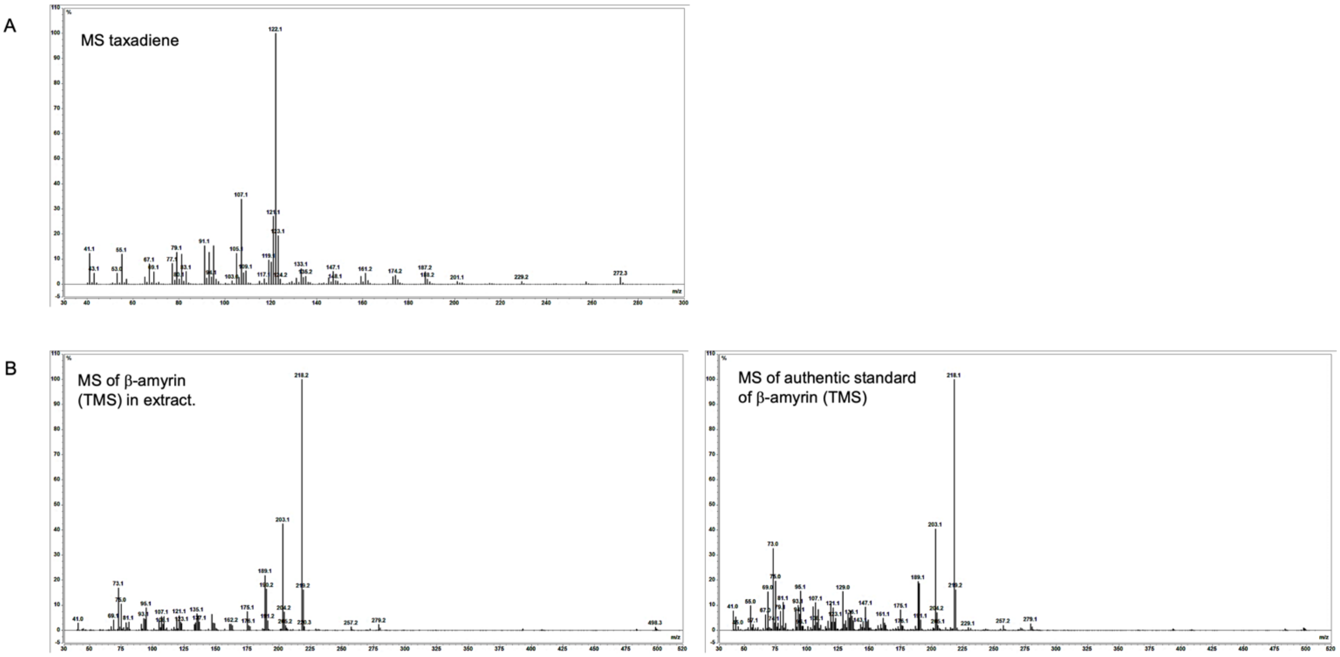


**Figure S3. Mass spectra of taxadiene and trimethylsilyl-derivatized β-amyrin identified in *Marchantia polymorpha* extracts.**

(A) Mass spectrum of taxadiene detected in *Marchantia polymorpha* terpene extracts. The molecular ion (M+) at 272 and fragment ion at 122 are characteristic of taxadiene. (B) Mass spectrum of trimethylsilyl-derivatized β-amyrin detected in *M. polymorpha* terpene extracts (left) compared to the authentic β-amyrin standard (right).

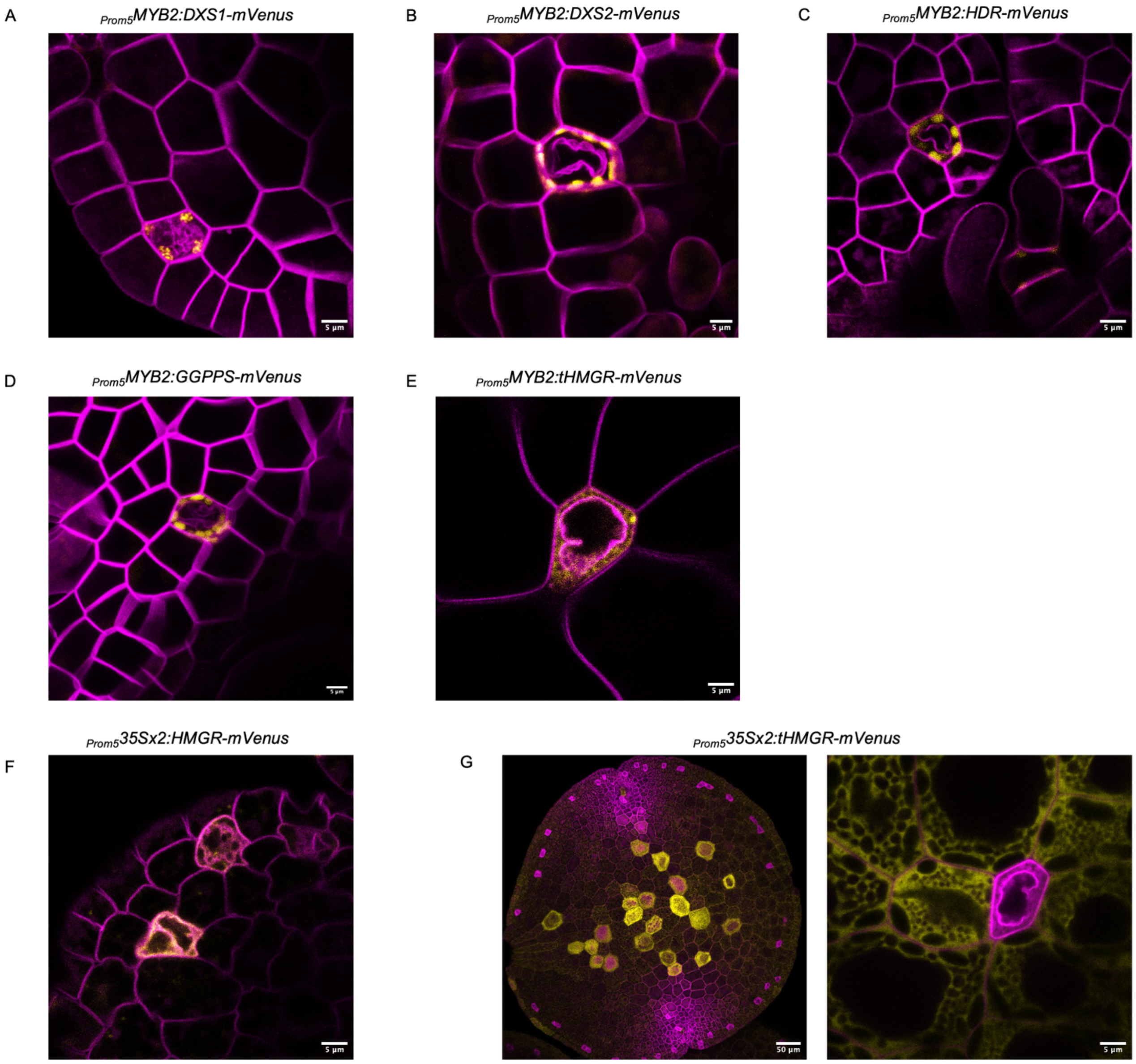


**Figure S4. Subcellular localization of precursor enzymes expressed under alternative promoters.**
Subcellular localization of *Marchantia* mVenus-tagged fusion proteins driven by the oil body-specific $_{Prom5}$*MYB2* promoter for (A) DXS1, (B) DXS2, (C) HDR, (D) GGPPS, and (E) tHMGR (scale bars: 5 µm). Subcellular localization of HMGR (F) and tHMGR (G) fusion proteins expressed under the *2x35S* promoter. For $_{Prom5}$*2x35S:tHMGR-mVenus* (G), a broader view of gemmae is shown (left panel; scale bar: 50 µm), along with a higher-magnification image (right panel; scale bar: 5 µm). Cell boundaries are marked by mScarlet fluorescence (purple).

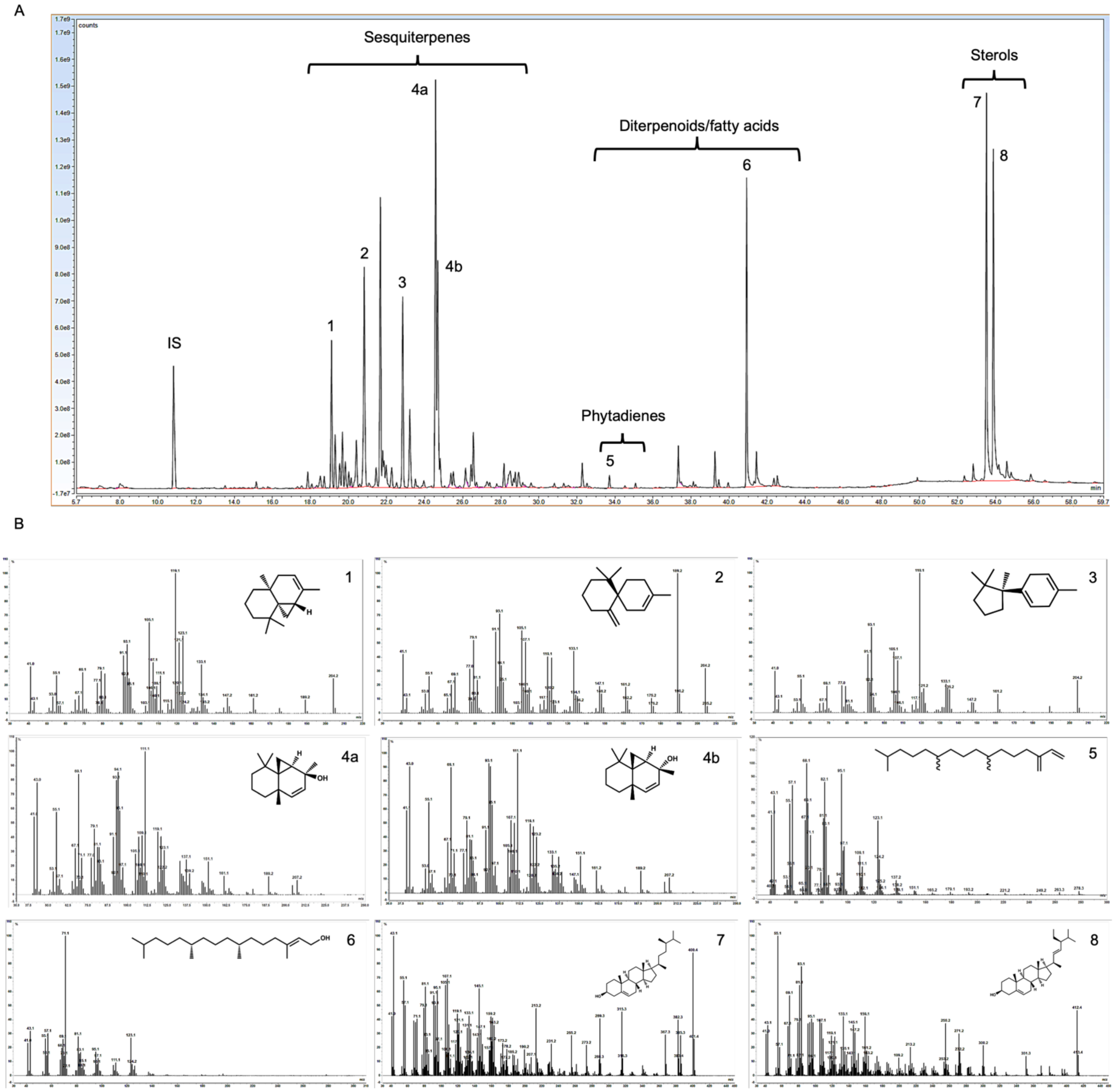


**Figure S5. Total ion chromatogram (TIC) and mass spectra of terpenes and phytosterols detected in *Marchantia polymorpha* extracts.**

(A) TIC of a methanol extract followed by hexane extraction from a 2-month-old, non-axenic *Marchantia polymorpha* culture. The selected peaks were tentatively identified as cis-thujopsene (1), β-chamigrene (2), γ-cuprenene (3), thujopsan-2α-ol (4a), thujopsan-2β-ol (4b), neophytadiene (5), phytol (6), campesterol (7), and stigmasterol (8). IS denotes the internal standard used for quantification. (B) Mass spectra of the labeled peaks in (A) with corresponding chemical structures of the putative compounds. Sesquiterpenes (1–4b) and diterpenes/fatty acids (5–6) were tentatively identified using Kovats retention indices and mass spectral data, while phytosterols (7 and 8) were identified based on their characteristic mass spectra.

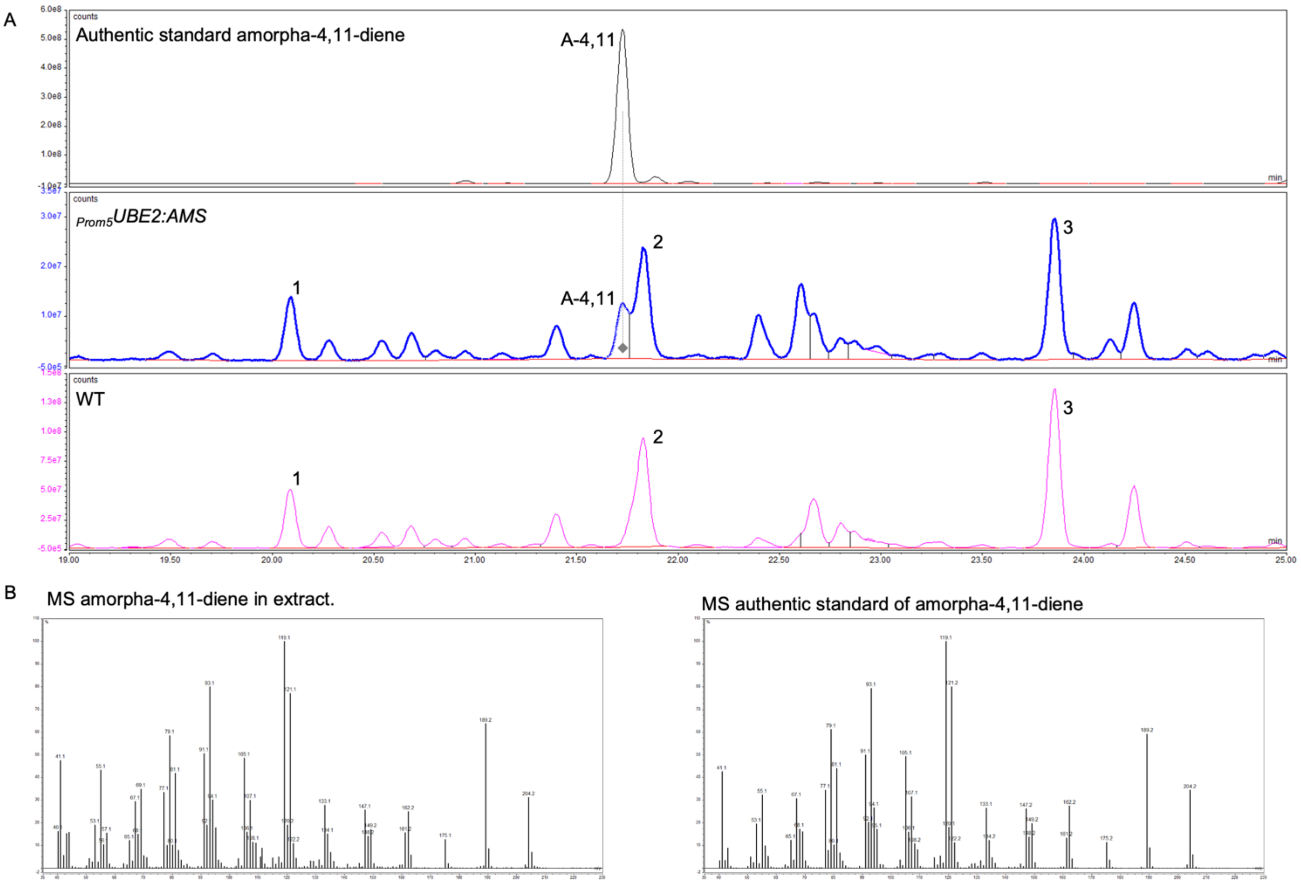


**Figure S6. Identification of amorpha-4,11-diene in *Marchantia polymorpha* expressing *Artemisia annua* AMS under $_{Prom5}$*UBE2*.**

(A) TICs of terpene extracts from WT *Marchantia polymorpha* (lower chromatogram), plants expressing $_{Prom5}$*UBE2:AMS* (middle chromatogram), and the authentic standard of amorpha-4,11-diene (upper chromatogram). A new peak labeled A-4,11 appears in the middle chromatogram, matching the retention time of the authentic standard. Peaks labeled 1, 2, and 3 correspond to the endogenous sesquiterpenes cis-thujopsene, β-chamigrene, and γ-cuprenene, respectively. (B) Mass spectra of amorpha-4,11-diene in the extract (left) and the authentic standard (right), demonstrating matching fragmentation patterns.

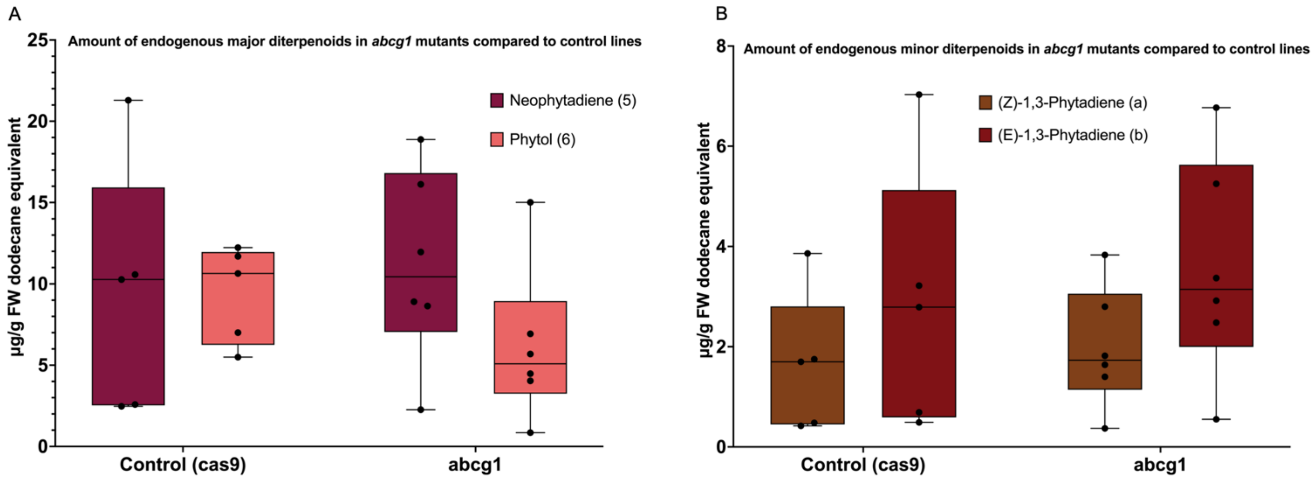


**Figure S7. Quantification of phytol and its derivatives in *abcg1* mutant and *Cas9* control lines.** **(**A) Levels of major fatty acids/diterpenoids neophytadiene (5) and phytol (6), in Cas9 controls (n=5) and abcg1 mutant lines (n=6). (B) Quantification of the minor fatty acids/diterpenoids (*Z*)-1,3-phytadiene (a) and (*E*)-1,3-phytadiene (b) in Cas9 controls (n=5) and abcg1 mutant lines (n=6). Box plots display individual data points for each compound, with bars representing the median and interquartile range.

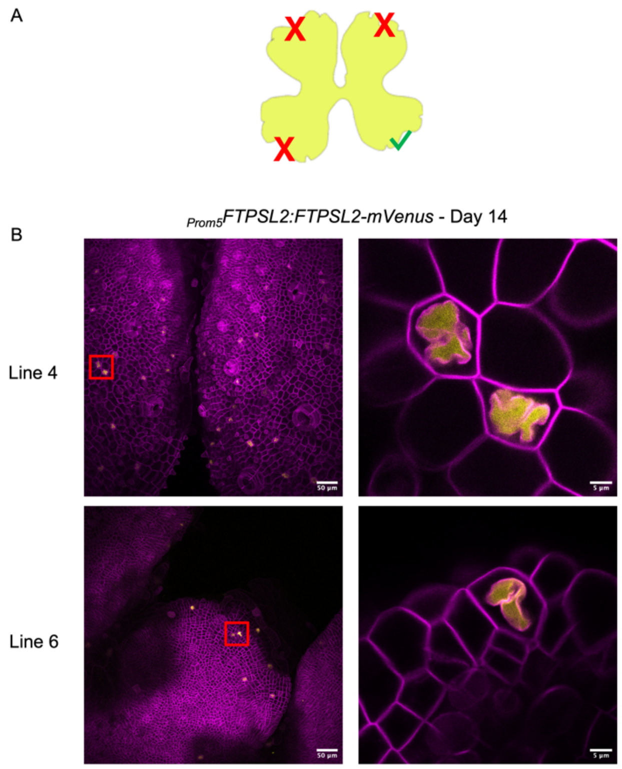


**Figure S8. Localization of the endogenous sesquiterpene synthase FTPSL2 driven by its own promoter.**

(A). Illustration of a 14-day-old *Marchantia* thallus (adapted from Marchantia.info) showing that the mVenus signal was detected in only one of the four thallus lobes (green checkmark) and absent in the others (red X). (B) Confocal imaging of the translational reporter $_{Prom5}$*FTPSL2:FTPSL2-mVenus* in two independent 14-day-old plants (lines 4 and 6). Left panels show the meristematic region (scale bar: 50 µm), with red squares indicating the areas magnified in the right panels (scale bar: 5 µm).
